# Patterns of species richness and endemism in bee and plant communities in California

**DOI:** 10.64898/2026.08.06.743382

**Authors:** Gilbert F. Alarcon-Cruz, Sarah Jacobs, Bruce G. Baldwin, Katja C. Seltmann, Gretchen Le Buhn

## Abstract

Understanding the spatial distribution of species and their patterns of endemism is necessary for establishing effective conservation priorities. Despite the vital pollination services bees provide, California’s native bee distribution patterns remain largely unexplored relative to plants and butterflies. We analyzed bee species richness and endemism across California and their concordance with plant distributions. Richness was high across areas of the California Floristic Province, including the Sierra Nevada, San Francisco Bay Area and Central Coast, South Coast Ranges, and the Transverse and Peninsular ranges. Bee endemism was more localized, concentrated in the San Joaquin Valley, eastern Sierra Nevada and adjacent Great Basin, Sierra Nevada foothills, and California deserts. Because richness and endemism appear to operate at different spatial scales and likely respond to different environmental drivers, effective conservation strategies must address both. Additionally, conservation plans that incorporate both plant and bee diversity are needed to achieve more comprehensive biodiversity protection.

## Introduction

Native bees, including both solitary and social species, are locally and globally important providers of pollination services that are critical to the reproduction of many plant species in both natural and agricultural systems^1^. However, a multitude of factors including habitat loss^2^, disease^3^, urbanization^4,5^, and introduced species^6–10^ have all contributed to a decline in bee species^11,12^. Recent analyses specific to North America identify pesticides and land use change as the primary drivers of bee decline, while invasive species effects appear less pronounced than in other regions^13^. Despite their importance, bee distribution patterns remain surprisingly understudied^14–17^ and little is known about the spatial patterns of richness and endemism of these taxa^2,18^. Understanding the current patterns of bee distribution will allow us to recognize when and how geographic ranges are changing and can help guide development of conservation priorities, especially as the climate changes^19^.

Spatial specimen-based occurrence data are useful to analyze patterns of species distributions across landscapes, and for identifying areas of low and high biodiversity^20,21^. Recently, there has been an increase in the availability of digital biodiversity data, from both museum and herbarium collections^22–24^ and observations from citizen science projects^25^ that enhance the ability to investigate the biodiversity of many organisms including bees^26–28^. Working with specimen-based spatial data remains challenging as many spatial data sets have important geographic gaps in the available records^29–31^. These gaps can arise from biased collecting^32,33^, which can occur even with an abundance of data^34^ or simply a lack of digitization or georeferencing of collections. Other challenges that may appear when working with digitally available data (i.e. occurrence data) include a lack of coordination between database systems and the increase in time required to analyze larger data sets^35^.

Concurrent with the availability of digitally available data, new tools for working with digital, specimen-based distributional data and addressing some of these issues have been developed^36^. This has led to identification of regions of high species richness and endemism for California vascular plants^37^ and for North American butterflies^38^. While specimen-based data have been used to illuminate the factors associated with increasing or decreasing abundance of bees^11,27,39^, North American patterns of distribution and endemism for solitary and social bees remain largely unexplored (but see Buckner et al.^14,15^).

There also remains a gap in research concerning the spatial relationships between bee and plant diversity at these coarser geographic scales. Both plant and bee distributions are a function of environmental and topographic factors^38,40–42^. The strong association of insect pollinators with angiosperms suggests that these two groups may have concordant biogeographic patterns^43–45^. The pattern for butterflies does not follow a strong association with angiosperms^38^. If there is concordance, identifying regions of shared high levels of endemism and richness in both plants and insects could provide an efficient template for conservation planning that would conserve more biodiversity^46^.

California is an ideal location to investigate the relationship between bees and plants due to its geography, high biodiversity, and extensive natural history collections^27^ and recent influential work on richness and endemism of California native plants^37^. California’s diverse geography encompasses both agricultural and natural landscapes, with approximately 1600 species of native bees^47,48^ and approximately 5,400 species of native plants^49^ making it a biodiversity hotspot of North America north of Mexico and an ideal location to investigate bee-plant relationships across varied land use contexts.

California can be divided into three biogeographic provinces (Figure 1, Table 1): the Desert Province, the Great Basin Province, and the California Floristic Province^50^. The Desert Province, occupying the southeastern portion of the state, is characterized by extreme aridity and high summer temperatures, with plant communities dominated by creosote bush (*Larrea tridentata*), cacti, and other drought-adapted shrubs. The Great Basin Province, covering the northeastern interior, is a cold semi-arid region of basin-and-range topography where sagebrush (*Artemisia* spp.) scrub and salt-tolerant vegetation predominate. The California Floristic Province is the largest and most botanically diverse province^49^, with a total of ca. 4,650 Californian species of vascular plants, >37% of which are endemic to the province^51^.

**Figure 1:**
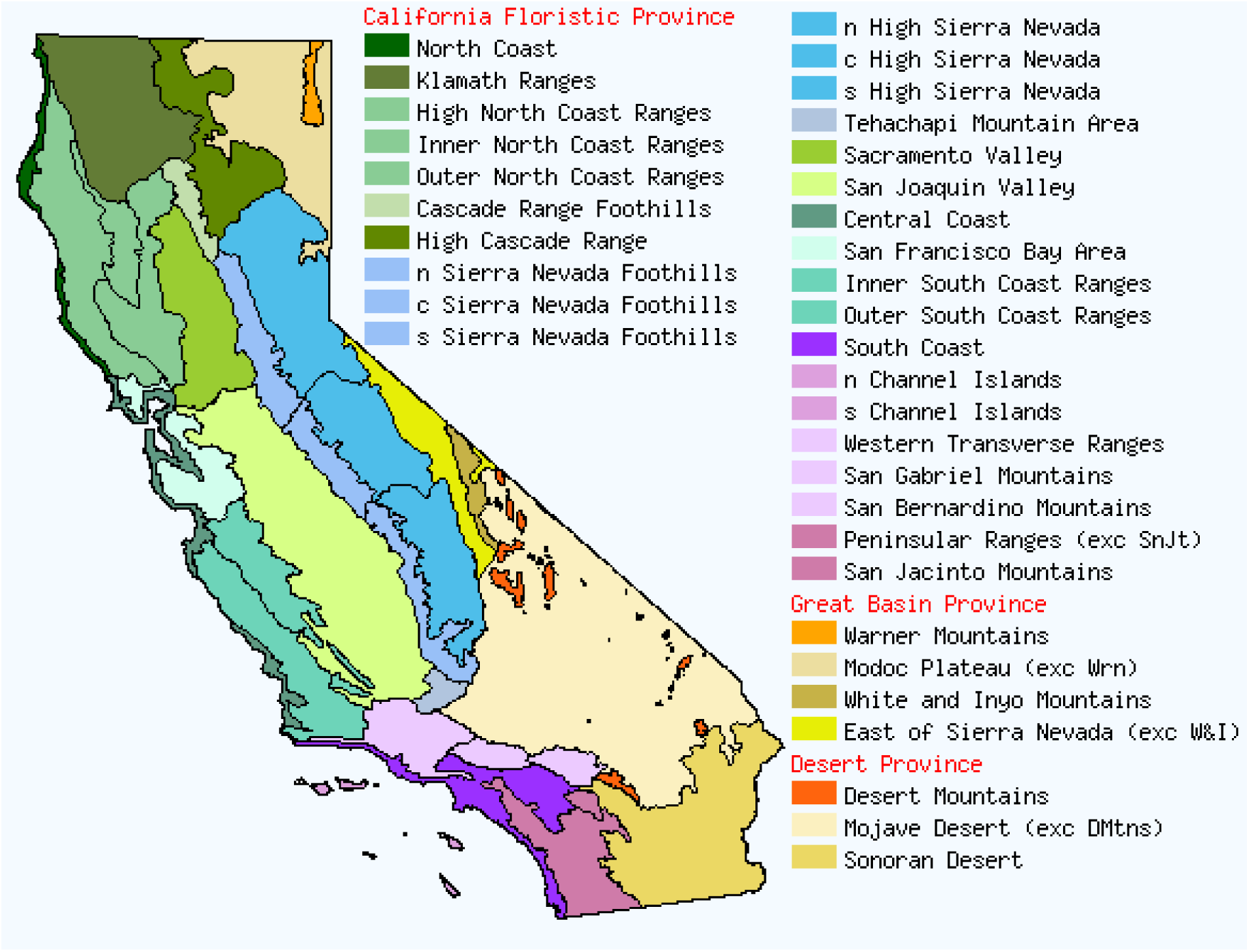
Geographic subdivisions of California, reproduced with permission of the Jepson Herbarium, University of California, Berkeley⁴⁹. Two-letter subdivision codes are given in Table 1 and are used in place of full names on all subsequent maps.

**Table 1.** Jepson subregions and their abbreviations.

| **Map label** | **Jepson subregion** |
| --- | --- |
| CCo | Central Coast |
| CaRF | Cascade Range Foothills |
| CaRH | High Cascade Ranges |
| ChI | Channel Islands |
| DMoj | Mojave Desert |
| DMtns | Desert Mountains |
| DSon | Sonoran Desert |
| KR | Klamath Ranges |
| MP exc Wrn | Modoc Plateau except Warner Mountains |
| NCo | North Coast |
| NCoR | North Coast Ranges |
| PR exc SnJt | Peninsular Ranges except San Jacinto Mountains |
| SCo | South Coast |
| SCoR | South Coast Ranges |
| SNE exc W&I | East of the Sierra Nevada except White and Inyo Mountains |
| SNF | Sierra Nevada Foothills |
| SNH | High Sierra Nevada |
| ScV | Sacramento Valley |
| SnFrB | San Francisco Bay Area |
| SnJV | San Joaquin Valley |
| SnJt | San Jacinto Mountains |
| TR | Transverse Ranges |
| Teh | Tehachapi Mountains Area |
| W&I | White and Inyo Mountains |
| Wrn | Warner Mountains |

Traditionally, bees have not been matched to ecoregions to the same extent as native plants. However, many bee genera have their greatest diversity in the mediterranean-like and montane climates of cismontane California (e.g., *Chelostoma*, *Dufourea*, and *Panurginus*)^52,53^, which broadly corresponds to the California Floristic Province.

The availability of geo-referenced datasets of bee collections offers the opportunity to do the first examination of patterns of richness and endemism of California bee species and the concordance of California bee and plant distribution patterns based on equal area spatial units. In this paper, we ask: What are the patterns of richness and endemism of bee species across California and do those patterns correspond to the patterns of angiosperm richness and endemism? To answer these questions, we followed the methods used by Baldwin et al.^37^ to find areas of endemism and species richness and expand that work by comparing the distributions of solitary and social bees and plants. Exploring these opportunities will help deepen the understanding of solitary bee and plant spatial distribution patterns in California and provide a template for other regions.

## Results

For ease of comparison, we present both the bee and angiosperm analyses side-by-side in the figures. Here, we focus on interpreting the results of the bee analysis and similarities and differences between the bee and plant distribution results. We do not discuss the specifics of the plant distribution results as they have already been presented in Baldwin et al.^37^.

### Data validation

From the bee occurrence data, Biodiverse generated results in 1,694 grid cells measuring 15 km × 15 km. The number of occurrences per cell ranged from 1 to 43,478 with a median of 44. Using the angiosperm data, Biodiverse generated results in 1,959 grid cells. The number of occurrences per cell ranged from 1 to 11,854 with a median of 330. In both bees (Figure 2a, 2b, and 2c) and plants^37^ (Figure 3a, 3b, and 3c), scatterplots showed a strong positive linear relationship between the number of occurrences and species richness (bees: slope = 0.664, R² = 0.881, p < 0.001). While weighted endemism (WE) and CWE) also showed significant positive relationships with occurrence counts in bees, the effect was substantially weaker (CWE: slope = 0.140, R² = 0.105, p < 0.001; Figure 2c), similar to the pattern observed in angiosperms^37^ (Figure 3c).

**Figure 2:**
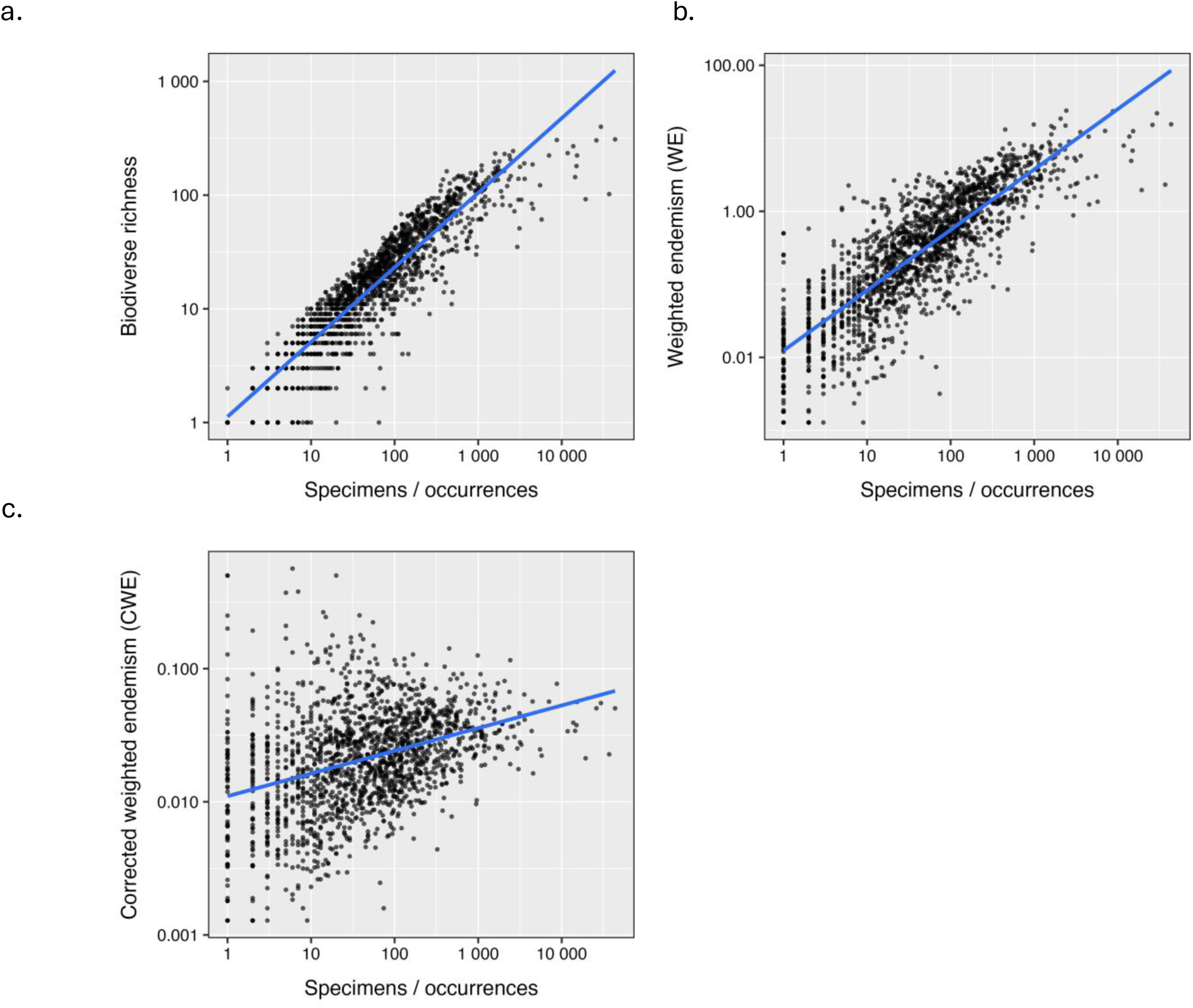
Relationship between the number of bee occurrence records per 15 km × 15 km grid cell and (a) species richness, (b) weighted endemism, and (c) corrected weighted endemism, across 1,694 cells. Lines are ordinary least-squares fits. Richness increases strongly with sampling effort (slope = 0.664, R² = 0.881, p < 0.001), whereas the relationship with corrected weighted endemism is much weaker (slope = 0.140, R² = 0.105, p < 0.001).

**Figure 3:**
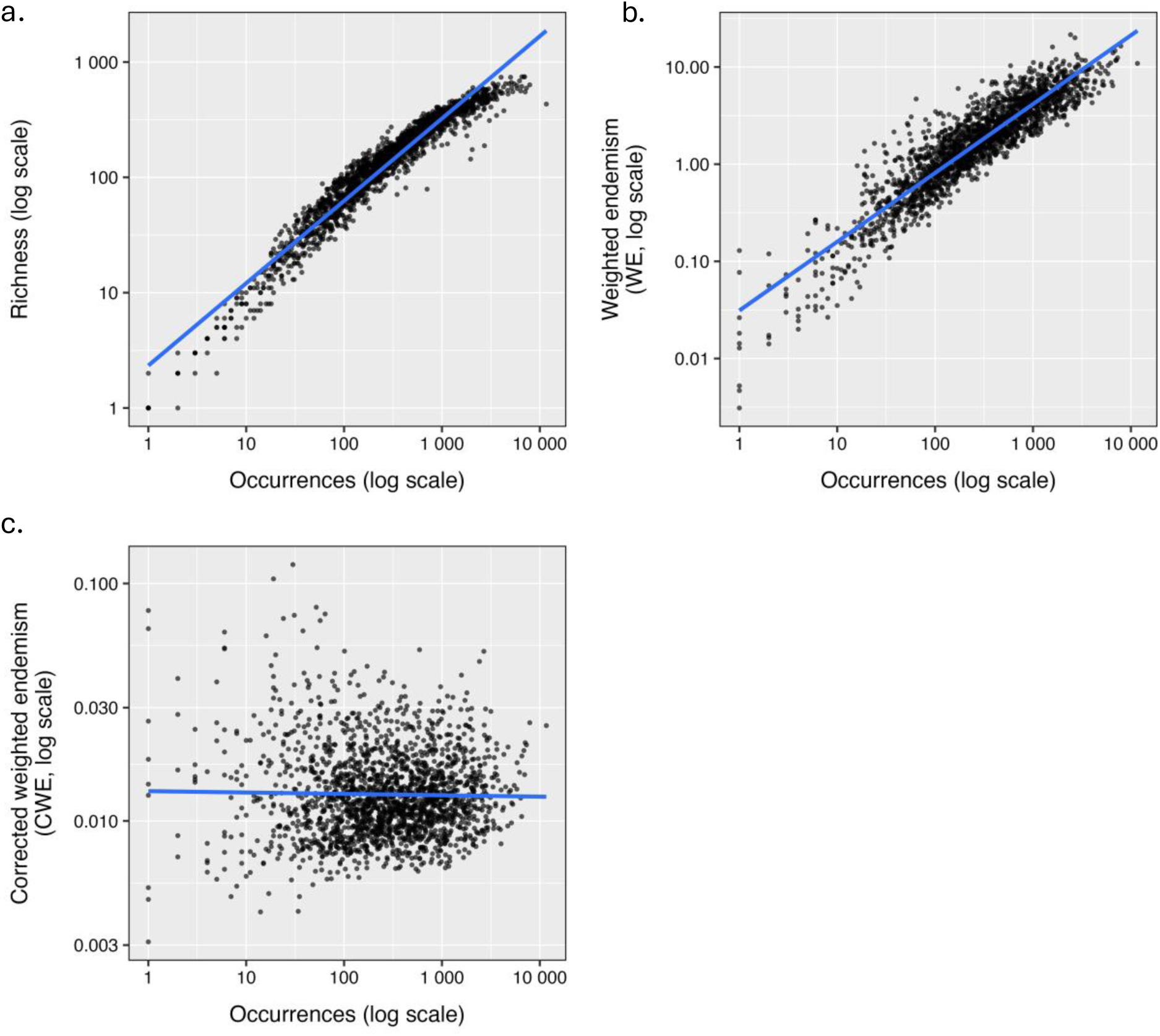
Relationship between the number of angiosperm occurrence records per 15 km × 15 km grid cell and (a) species richness, (b) weighted endemism, and (c) corrected weighted endemism, across 1,959 cells. Lines are ordinary least-squares fits. As in bees, richness scales strongly with sampling effort while corrected weighted endemism does not.

Maps of redundancy reveal variation in sampling effort for bees across California (Figure 4a). Most cells were reasonably well sampled for bees (redundancy > 0.54^54^). However, there were some (∼200) cells with no sampling (indicated in white on the figure), particularly in the northern regions and the desert southeast. In contrast, angiosperm sampling (Figure 4b) was more evenly distributed, with fewer cells lacking sampling effort but also fewer cells with extensive sampling. Comparison of observed richness and Chao1-estimated richness across 1,695 grid cells showed that, although observed richness underestimated total richness as expected, the two metrics identified broadly similar sets of species-rich cells. Rank correlation between the two measures was high (Spearman ρ = 0.975; Figure S1), and overlap in the top-ranked cells was 85% for the top 10% of cells, 88% for the top 20%, and 91% for the top 30%, with corresponding Jaccard similarities of 0.73, 0.79, and 0.83 (Table S7). Median estimated completeness was 0.65 among cells with at least ten observed species, and varied little across the upper richness gradient (0.62–0.71 across richness deciles 5–10), indicating that undersampling is broadly comparable among cells rather than systematically concentrated in particular parts of the richness distribution. Completeness values approaching 1.0 in cells with very few records are an artifact of the estimator rather than evidence of complete inventory: 393 cells (23%) contain no doubleton species, so the Chao1 correction term approaches zero and undersampling cannot be detected. Together these results indicate that use of Chao1 would not substantially change the broad spatial pattern of cell ranking, although they confirm that a substantial fraction of bee species in any given cell remains undetected.

**Figure 4.**
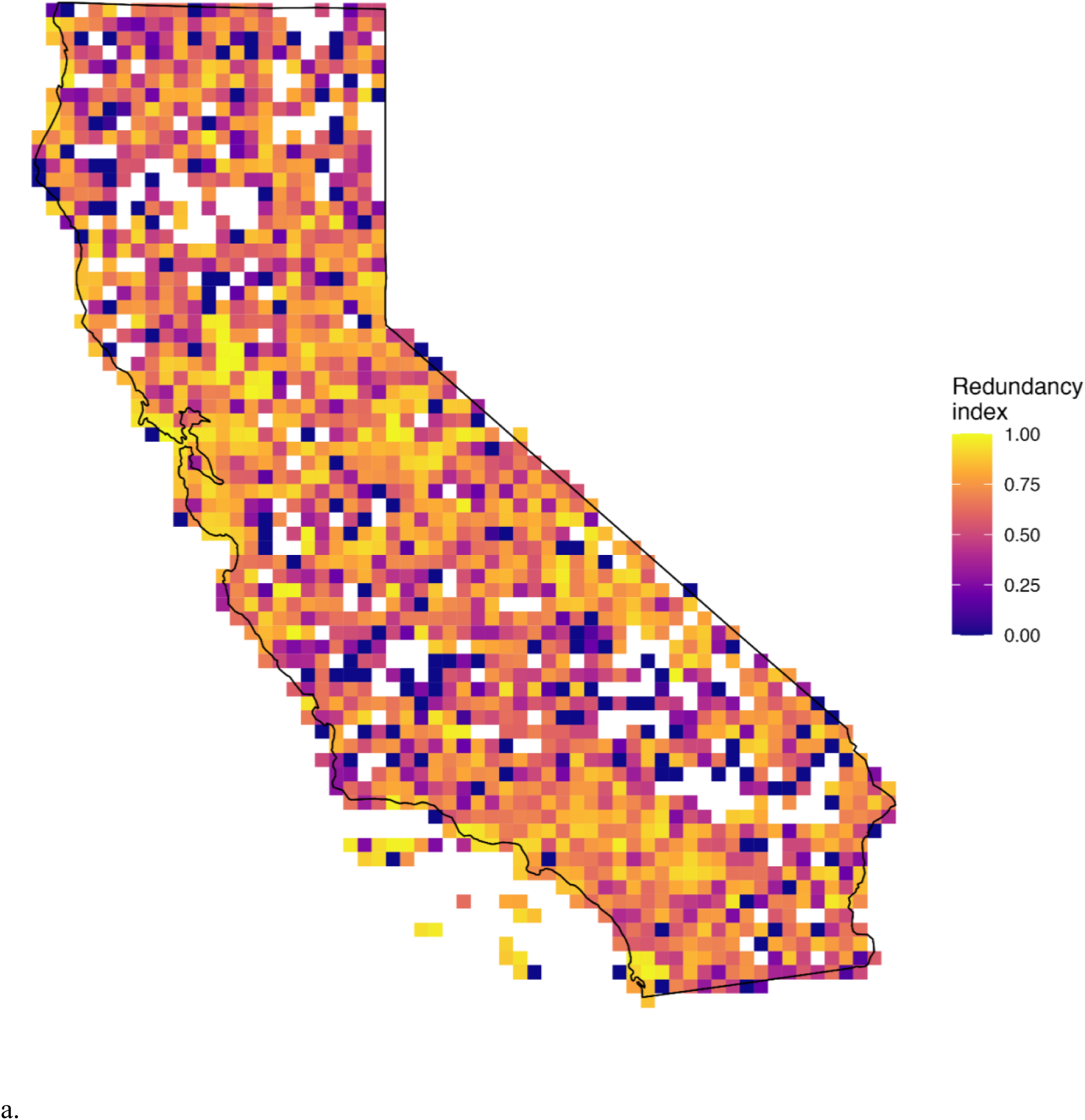
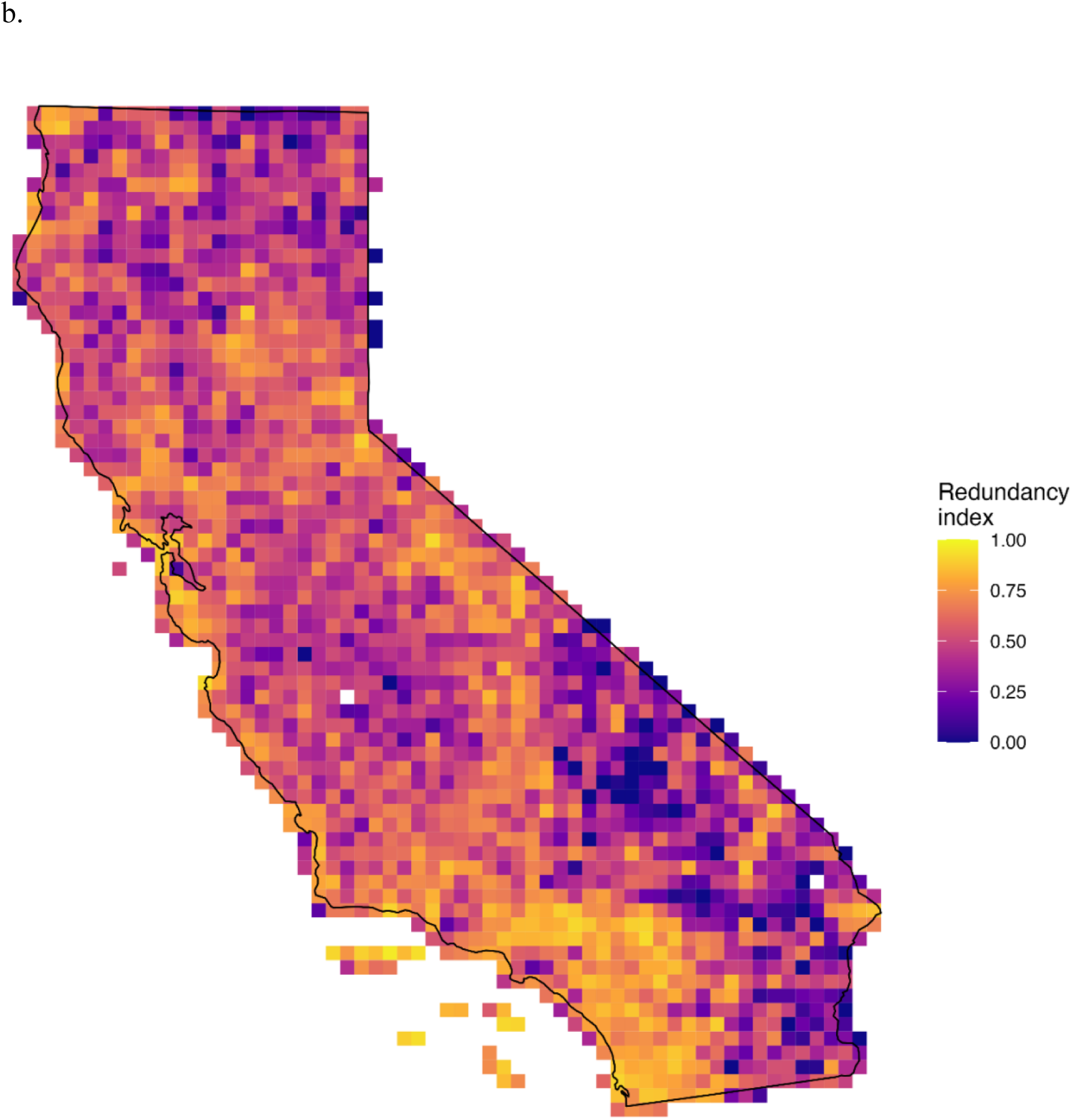
a. Sampling redundancy per 15 km × 15 km grid cell for (a) bees and (b) angiosperms. Redundancy is calculated from the number of records relative to the number of species recorded in each cell; values approaching 1 indicate cells whose species list is supported by repeated sampling, and low values indicate cells whose apparent richness rests on few records. Cells shown in white contain no records. Approximately 200 cells were unsampled for bees, concentrated in the northern interior and the southeastern deserts; angiosperm sampling was more evenly distributed. Jepson subdivision codes are given in Table 1.

### Observed patterns of bee and angiosperm species richness and endemism

Maps of observed bee species richness (Figure 5a) indicate the highest richness across the Sierra Nevada (SN), the eastern Transverse Ranges (TR), specifically the San Gabriel and San Bernardino mountains, and the Peninsular Ranges including San Jacinto Mountains (PR exc SnJt, SnJt) of southwestern California, the South Coast (SCo) and South Coast Ranges (SCoR) in the vicinity of Pinnacles National Park, and the San Francisco Bay Area (SnFrB) and adjacent Central Coast (CCo). In comparison, angiosperms also had large areas of high species richness across the Sierra Nevada (SN) and along the Pacific coast, including the San Francisco Bay Area (SnFrB), the South Coast Ranges (SCoR), and the Transverse and Peninsular ranges (TR, PR exc SnJt, SnJt) (Figure 5b). Unlike bees, there is also high angiosperm species richness in the northwestern part of the state, including areas of the North Coast (NCo), North Coast Ranges (NCoR), and Klamath Ranges (KR) of California.

**Figure 5.**
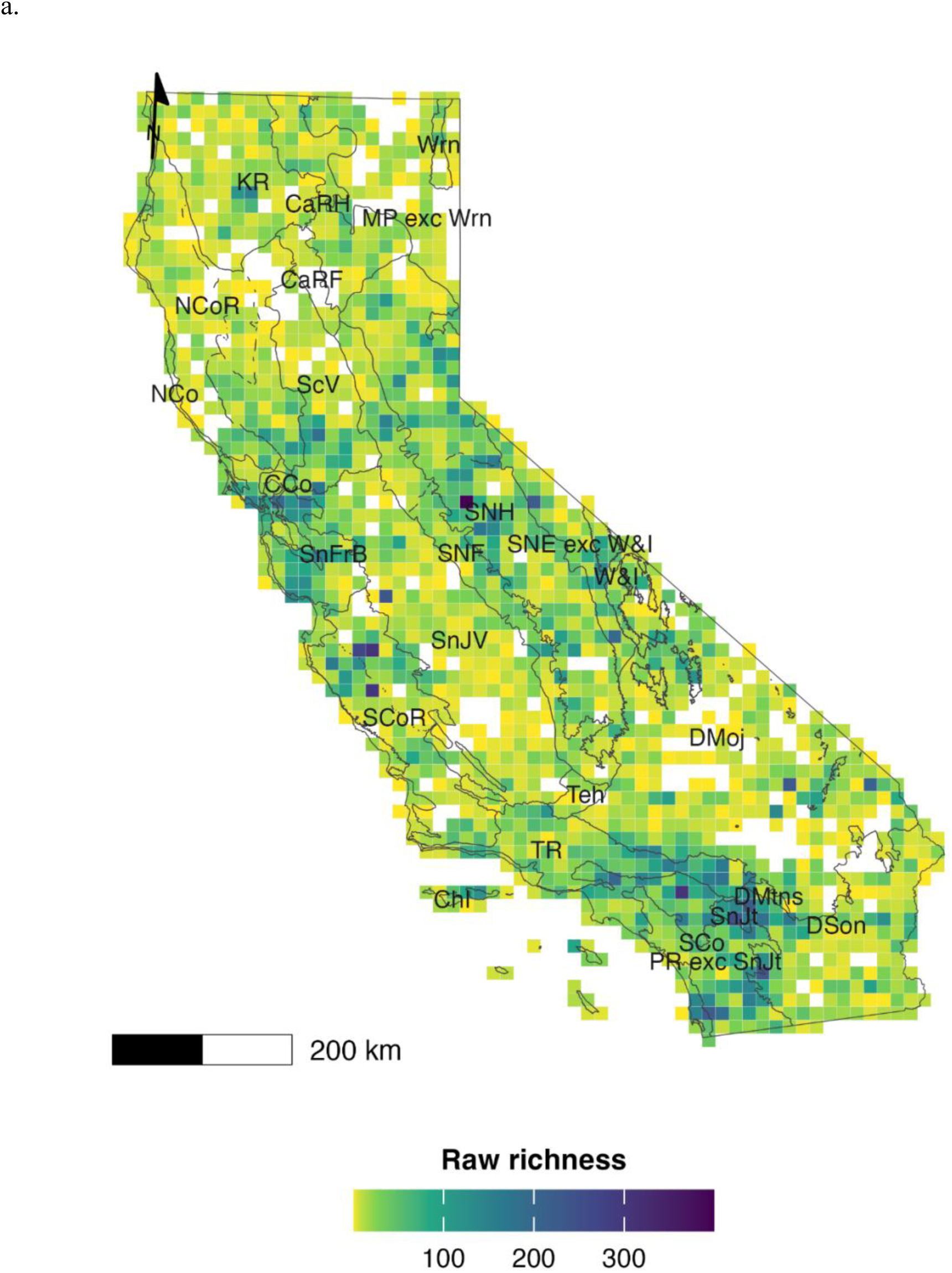
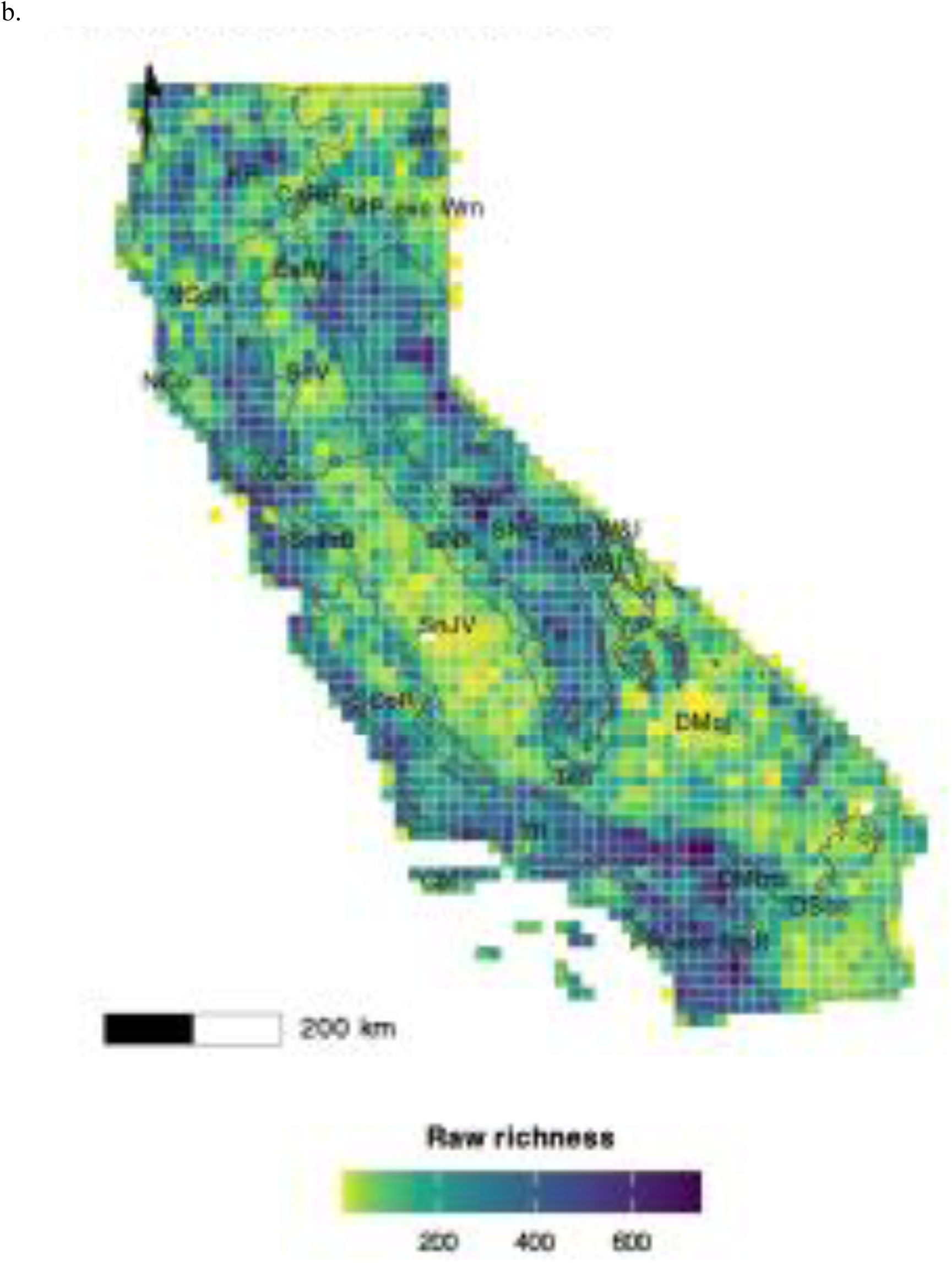
Observed species richness per 15 km × 15 km equal-area grid cell for (a) bees and (b) native angiosperms, calculated in Biodiverse v4.3. Cells with no recorded value are shown in grey. Grey outlines and two-letter codes mark Jepson geographic subdivisions (Table 1). Coordinate reference system: California Albers (EPSG:3310).

The distribution of areas with high CWE for bees, which highlights areas with a higher concentration of range-restricted species relative to their overall diversity (Figure 6a), are more restricted. CWE for bees is locally high within areas of the Mojave and Sonoran Desert (DMoj and DSon), especially in the west, Central Sierra Nevada Foothills (SNF), San Joaquin Valley (SnJV), High Cascade Ranges (CaRH), the Modoc Plateau excluding the Warner Mountains (MP exc Wrn), and East of Sierra Nevada (SNE). Angiosperm CWE is high in the White and Inyo Mountains (W&I), elsewhere East of Sierra Nevada (SNE exc W&I) and in some parts of other regions, including the eastern Mojave Desert (DMoj) and Modoc Plateau (MP) and Warner Mountains (Wrn), Klamath Ranges (KR), northern North Coast (NCo) and Channel Islands (ChI). Plant areas with high CWE are also locally restricted (Figure 6b).

**Figure 6:**
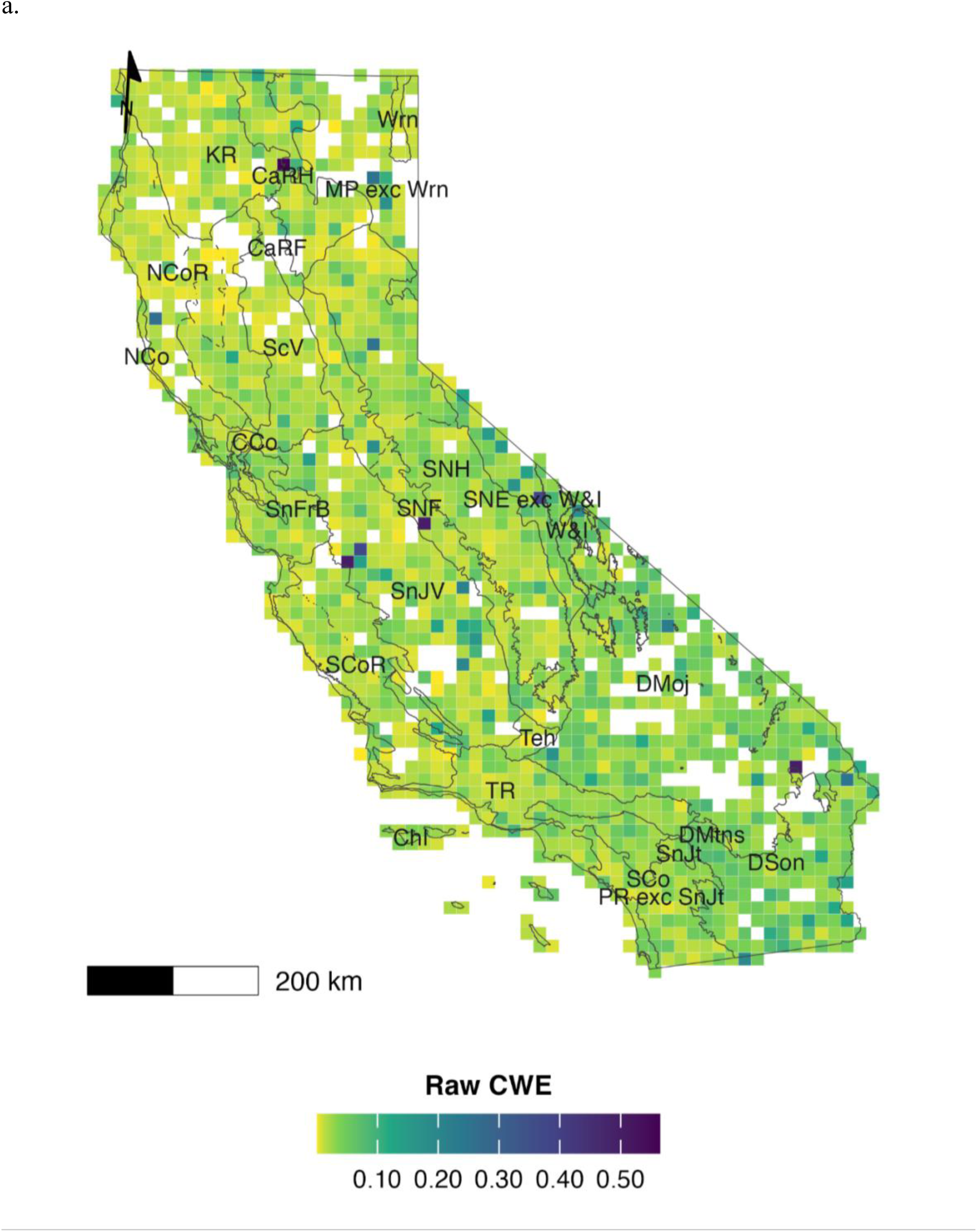
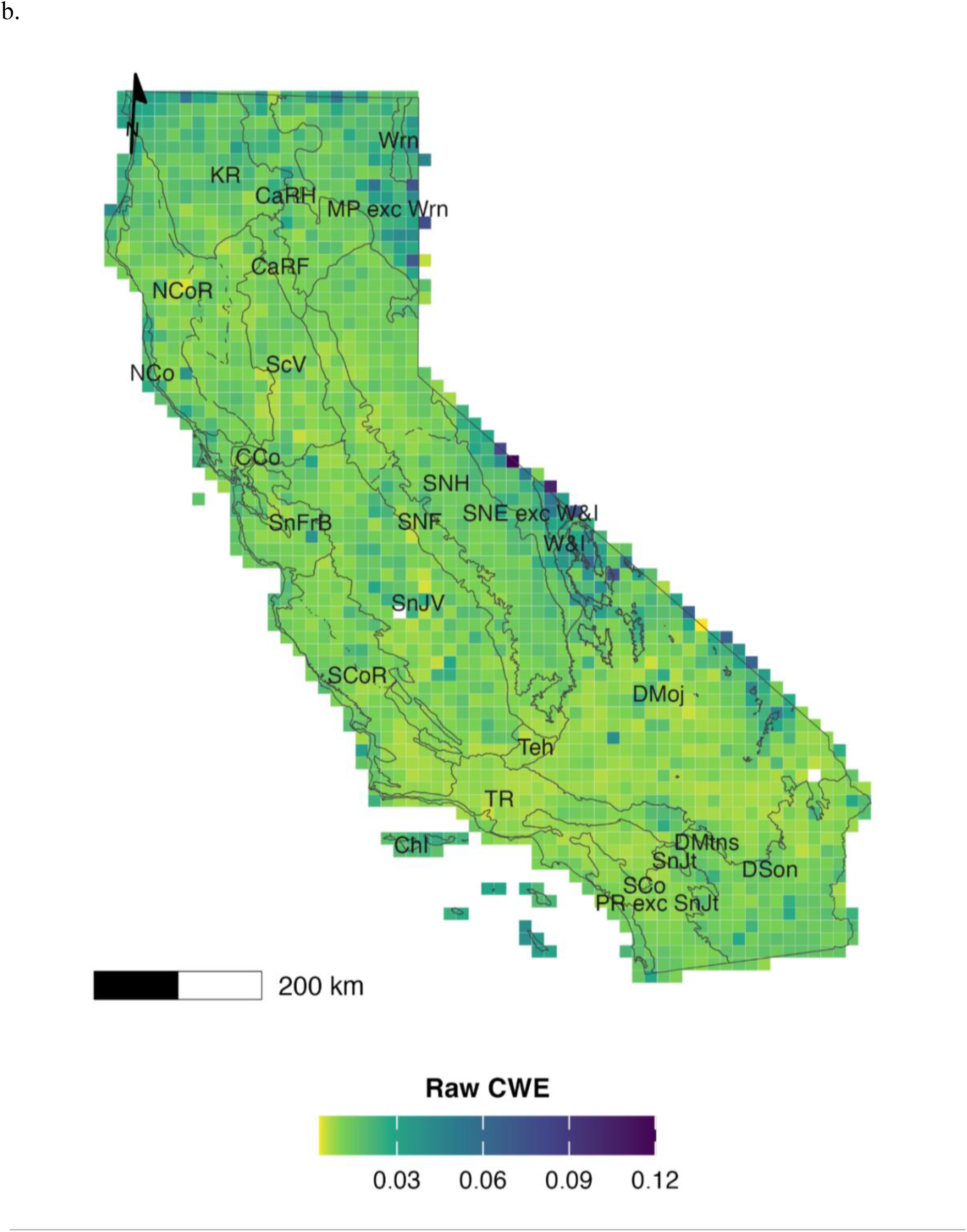
Corrected weighted endemism (CWE) per 15 km × 15 km equal-area grid cell for (a) bees and (b) native angiosperms, calculated in Biodiverse v4.3 as weighted endemism divided by cell species richness. CWE therefore expresses the average range- restrictedness of the species present rather than their number. epson subdivision codes are given in Table 1.

### Hotspot analysis of species richness and corrected weighted endemism

*Species richness.* A hotspot analysis of bee species richness (Figure 7a) indicated several hotspots with concentrated high species richness. One hotspot is made up of the San Francisco Bay Area (SnFrB), Central Coast (CCo), and southwestern Sacramento and northwestern San Joaquin valleys (ScV, SnJV). Another hotspot is made up of the Inner and northern Outer South Coast Ranges (SCoR). An additional hotspot is made up of the Central High Sierra Nevada and the adjacent southeastern Northern High Sierra Nevada (SNH), eastern Central Sierra Nevada Foothills (SNF), and the White and Inyo Mountains (W&I). The final hotspot is made up of the eastern Transverse Ranges (TR), specifically the San Gabriel and San Bernardino mountains, South Coast (SCo), and Peninsular Ranges including San Jacinto Mountains (PR exc SnJt, SnJt). Several small coldspots with concentrated low species richness are widely scattered, e.g., between the Modoc Plateau (MP), Northern High Sierra Nevada (SNH), San Joaquin Valley (SnJV), and Mojave Desert (DMoj).

**Figure 7:**
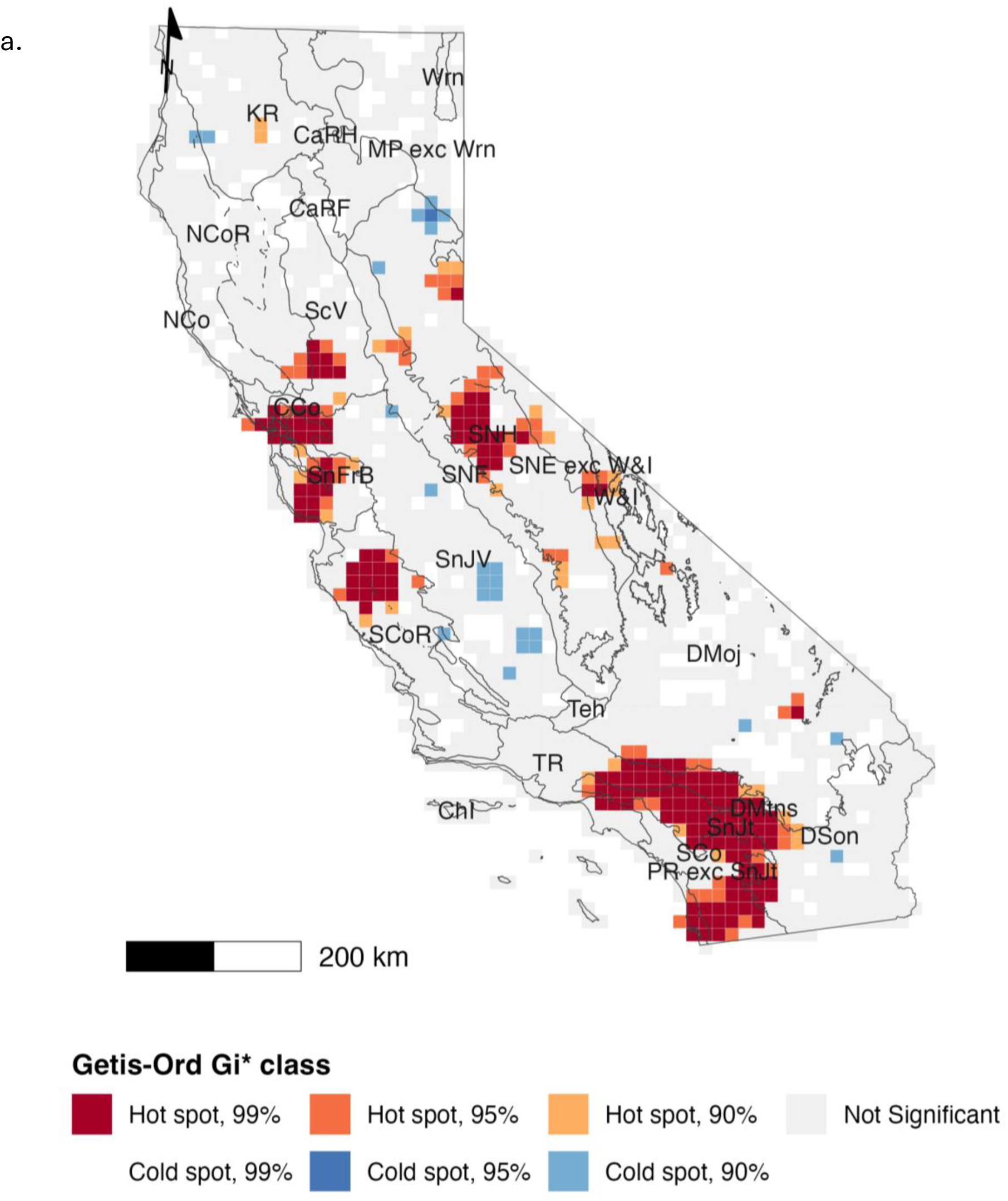
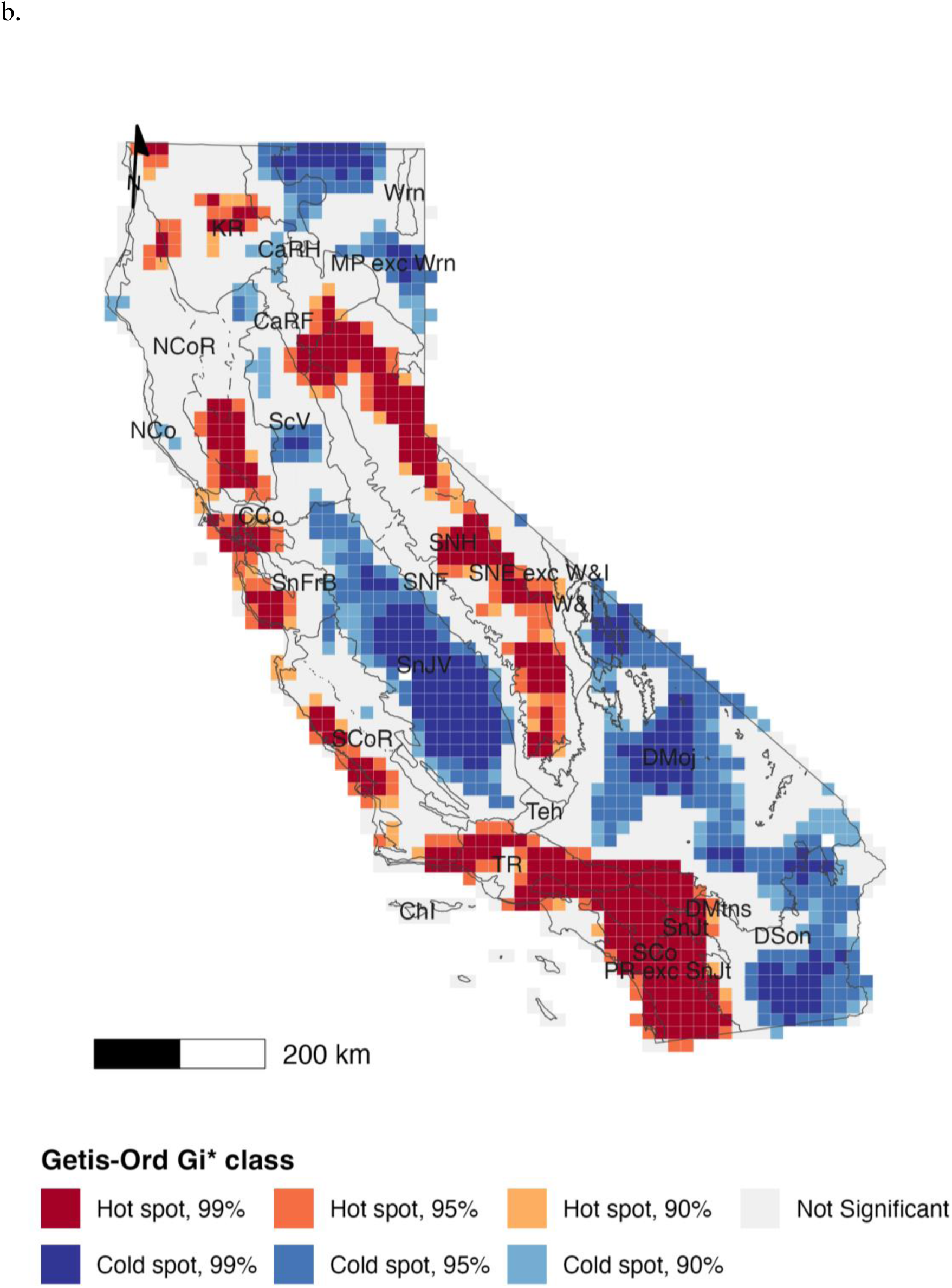
Getis-Ord Gi* hot spot analysis of species richness for (a) bees and (b) native angiosperms, using queen contiguity. Local Gi* statistics were computed for each 15 km × 15 km cell with a neighbour structure that includes the focal cell, under binary spatial weights. Two-sided p values derived from the Gi* z scores classify each cell as a hot spot or cold spot at the 90%, 95%, or 99% confidence level, or as not significant; p values are not adjusted for multiple comparisons. Jepson subdivision codes are given in Table 1. Coordinate reference system: California Albers (EPSG:3310).

A hotspot analysis of angiosperm species richness identified many more areas with distinct concentrations of high species richness, mostly in montane areas of the California Floristic Province (Figure 7b). The montane regions of the Klamath Ranges (KR), High Cascade Ranges (CaRH), Cascade Range Foothills (CaRF), High Sierra Nevada (SNH), Inner North Coast Ranges (NCoR), Outer South Coast Ranges (SCoR), Western Transverse Ranges (TR), and a large area encompassing the Transverse Ranges (TR, including the San Gabriel and San Bernardino mountains), Peninsular Ranges including San Jacinto Mountains (PR exc SnJt, SnJt), and the South Coast (SCo), and the foothill and plains regions of East of Sierra Nevada excluding White and Inyo Mountains (SNE exc W&I), San Francisco Bay Area (SnFrB), and Central Coast (CCo) were identified as hotspots with high species richness for angiosperms (Figure 7b). More striking for angiosperms are the extensive coldspots through the Sacramento and San Joaquin valleys (ScV, SnJV), the Mojave and Sonoran deserts (DMoj, DSon), and the High Cascade Ranges (CaRH) and Modoc Plateau (MP).

### Corrected weighted endemism

A hotspot analysis of bee CWE (Figure 8a) revealed smaller more widely distributed hotspots than richness analysis (Figure 7a). The largest hotspots are found in the western San Joaquin Valley and adjacent Inner South Coast Ranges (SnJV, SCoR), East of Sierra Nevada (SNE), Central Sierra Nevada Foothills (SNF), southern San Joaquin Valley (SnJV), eastern Klamath and adjacent High Cascade ranges (KR, CaRH), and several areas in the Mojave and Sonoran deserts (DMoj, DSon). The North Coast Ranges (NCoR) of northwest California comprise coldspots for bee corrected weighted endemism.

**Figure 8:**
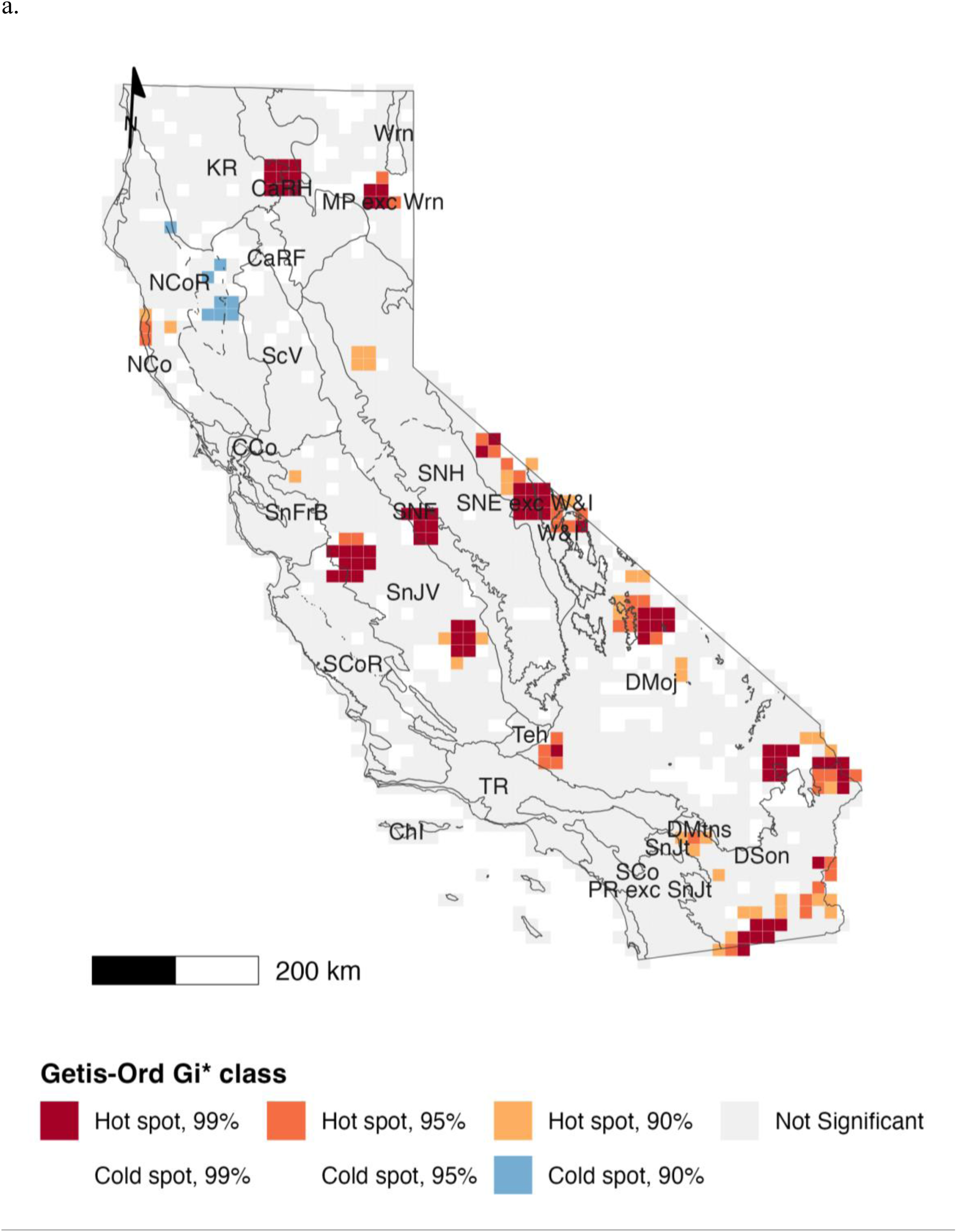
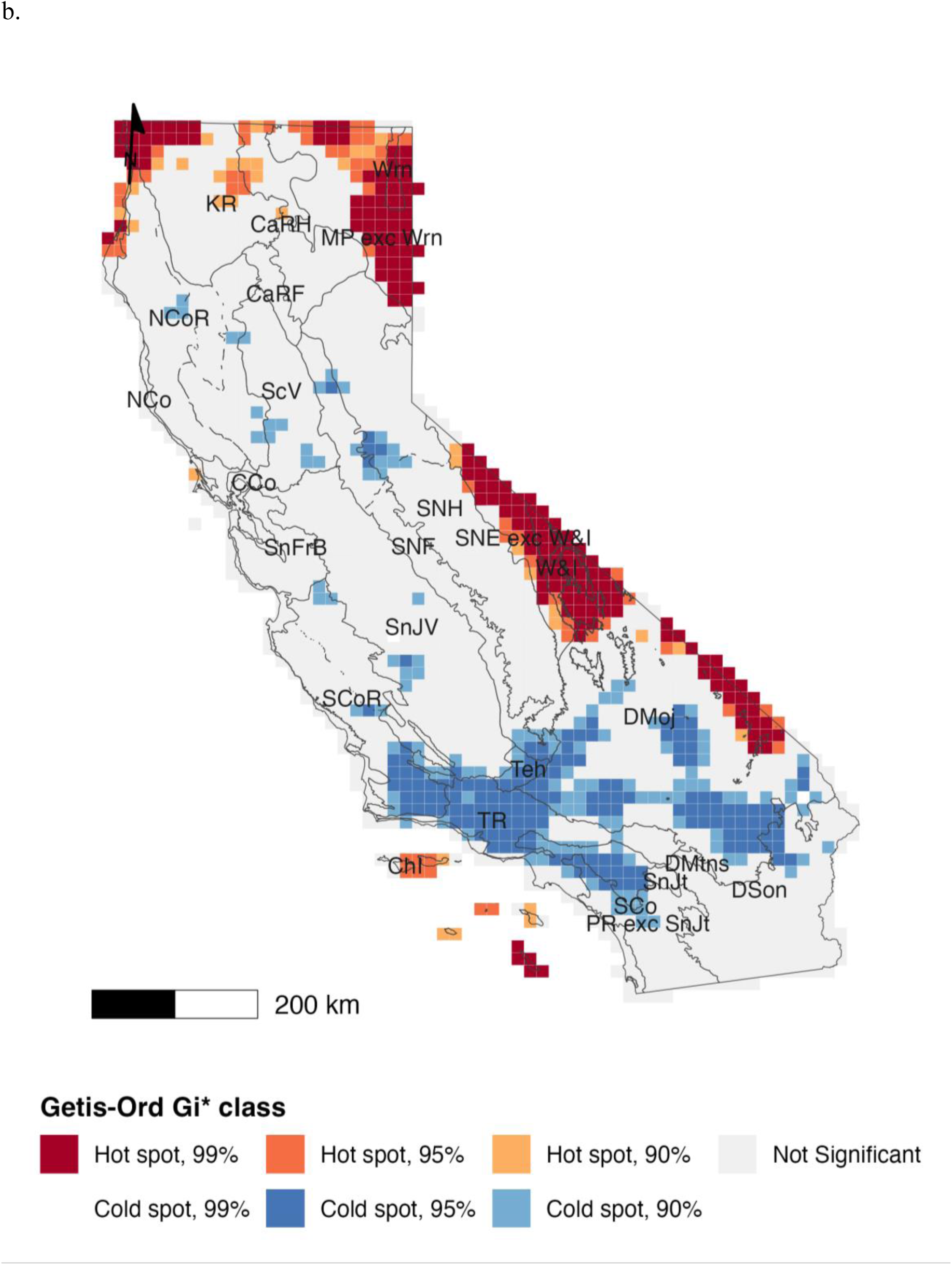
Getis-Ord Gi* hot spot analysis of corrected weighted endemism for (a) bees and (b) native angiosperms, using queen contiguity. Neighbour structure, weighting, significance thresholds, and projection follow Figure 7; p values are not adjusted for multiple comparisons. Jepson subdivision codes are given in Table 1.

A hotspot analysis of angiosperm corrected weighted endemism (Figure 8b) showed that hotspots are concentrated in northern and eastern California as well as the Channel Islands whereas the hotspots for angiosperm species richness are primarily in the Sierra Nevada and southern and western parts of the state. The areas with hotspots of angiosperm CWE include White and Inyo Mountains (W&I), East of Sierra Nevada (SNE) and in some parts of other regions, including the eastern Mojave Desert (DMoj), Modoc Plateau (MP), Warner Mountains (Wrn), and some parts of the Klamath and Cascade ranges (KR, CaRH), northern North Coast (NCo) and Channel Islands (ChI). There are extensive coldspots in the Transverse Ranges (TR, including the San Gabriel and San Bernardino mountains), Tehachapi Mountains (Teh), and northern Peninsular Ranges (PR) as well as areas of the central, western, and southern Mojave Desert (DMoj) and northern Sonoran Desert (DSon).

### Spatial autocorrelation in bee diversity patterns

Global spatial autocorrelation analyses revealed significant positive spatial clustering for both bee species richness and bee corrected weighted endemism, although the strength of autocorrelation differed markedly between metrics. Species richness showed moderate positive spatial autocorrelation, indicating broad spatial clustering of richness values among neighboring cells (Moran’s I = 0.300, *z* = 22.42, one-tailed *p* < 0.0001; permutation *P* = 0.0001). Corrected weighted endemism showed weaker, but still highly significant, positive spatial autocorrelation (Moran’s I = 0.089, *z* = 6.80, one-tailed *p* < 0.0001; permutation *P* = 0.0001). Thus, spatial autocorrelation in corrected weighted endemism was approximately threefold lower than that observed for species richness. The Moran scatterplot for richness (Figure 9a) shows a positive slope, consistent with moderate positive spatial autocorrelation broad spatial clustering of richness values. Cells with high richness tended to occur near other high-richness cells and low-richness cells tended to occur near low-richness neighbors. In contrast, the Moran scatterplot for endemism (Figure 9b) exhibits a shallow positive slope, indicating weak but significant spatial autocorrelation.

**Figure 9.**
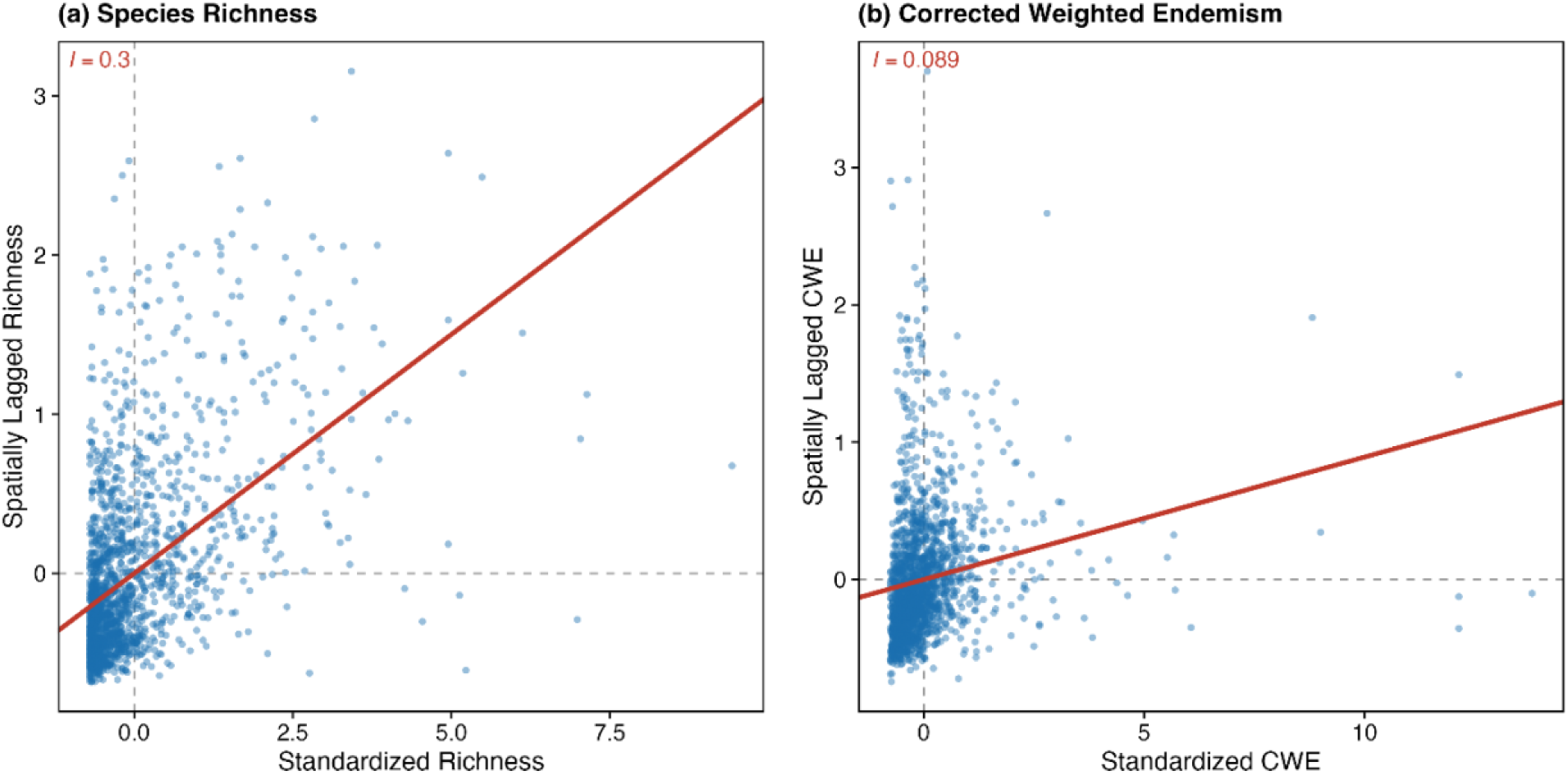
Moran scatterplots showing global spatial autocorrelation in (a) bee species richness and (b) bee corrected weighted endemism. Each point is one 15 km × 15 km grid cell, with the x-axis showing the standardized metric value and the y-axis its spatially lagged value — the weighted average across neighboring cells under a row-standardized queen-contiguity weights matrix. Points in the upper-right quadrant are high-value cells surrounded by high-value neighbors and points in the lower-left are low-value cells surrounded by low-value neighbors, both indicating positive spatial association; points in the upper-left and lower-right quadrants are spatial outliers. The red line is the ordinary least-squares fit, whose slope equals Moran’s I; the value is annotated on each panel.

### Comparison of bee and angiosperm patterns of species richness

Similarity between angiosperm and bee richness was largely confined to hotspots (Figure 10a). Shared richness hotspots (both taxa significantly high) cover approximately 9% of the analysis grid and were strongly concentrated in southern California, forming a nearly continuous block across the eastern Transverse Ranges (TR), the Peninsular Ranges, including the San Jacinto Mountains (PR exc SnJt, SnJt), the adjacent South Coast (SCo) and desert mountains (DMtns), and extending onto the western margins of the Sonoran and Mojave deserts (DSon, DMoj) where they abut those ranges. Roughly two-thirds of all shared hotspots occurred in this southern block. Elsewhere, shared hotspots were scattered rather than contiguous: along the crest of the High Sierra Nevada (SNH) and the adjacent area to the east (SNE exc W&I, W&I), distributed from the northern through the southern Sierra rather than in discrete northern, central, and southern clusters; and in the San Francisco Bay Area (SnFrB) with adjacent cells in the Central Coast (CCo) and northern South Coast Ranges (SCoR). Shared coldspots, by contrast, were rare (<1% of cells) and were nearly restricted to the southern San Joaquin Valley (SnJV), with isolated cells in the Mojave and Sonoran deserts. The interior lowlands and deserts (the Mojave Desert (DMoj), San Joaquin Valley (SnJV), Sonoran Desert (DSon), Modoc Plateau (MP), and Sacramento Valley (ScV)) were dominated by angiosperm-only coldspots, indicating significantly low plant richness where bee richness was not significantly low.

**Figure 10:**
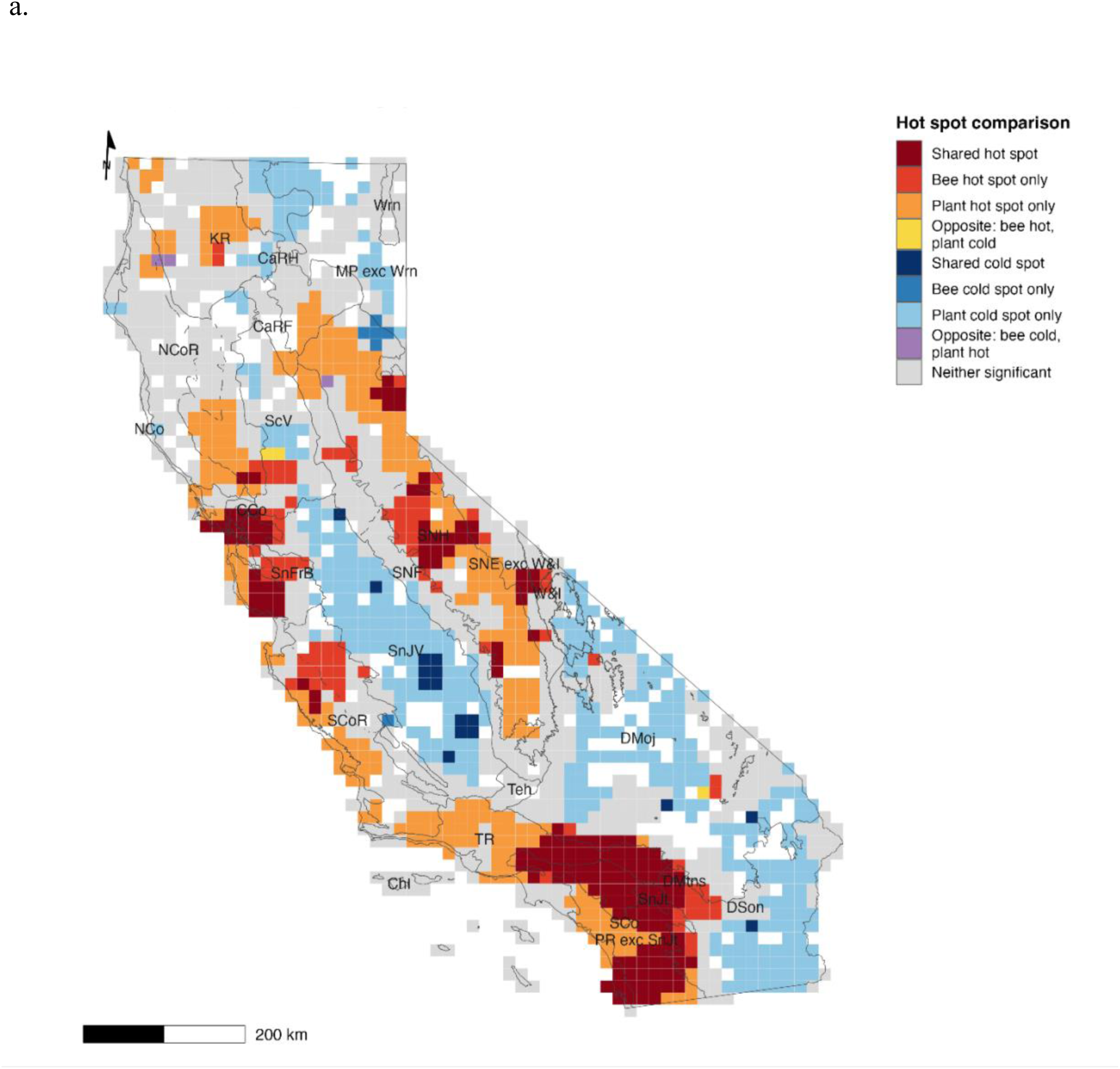
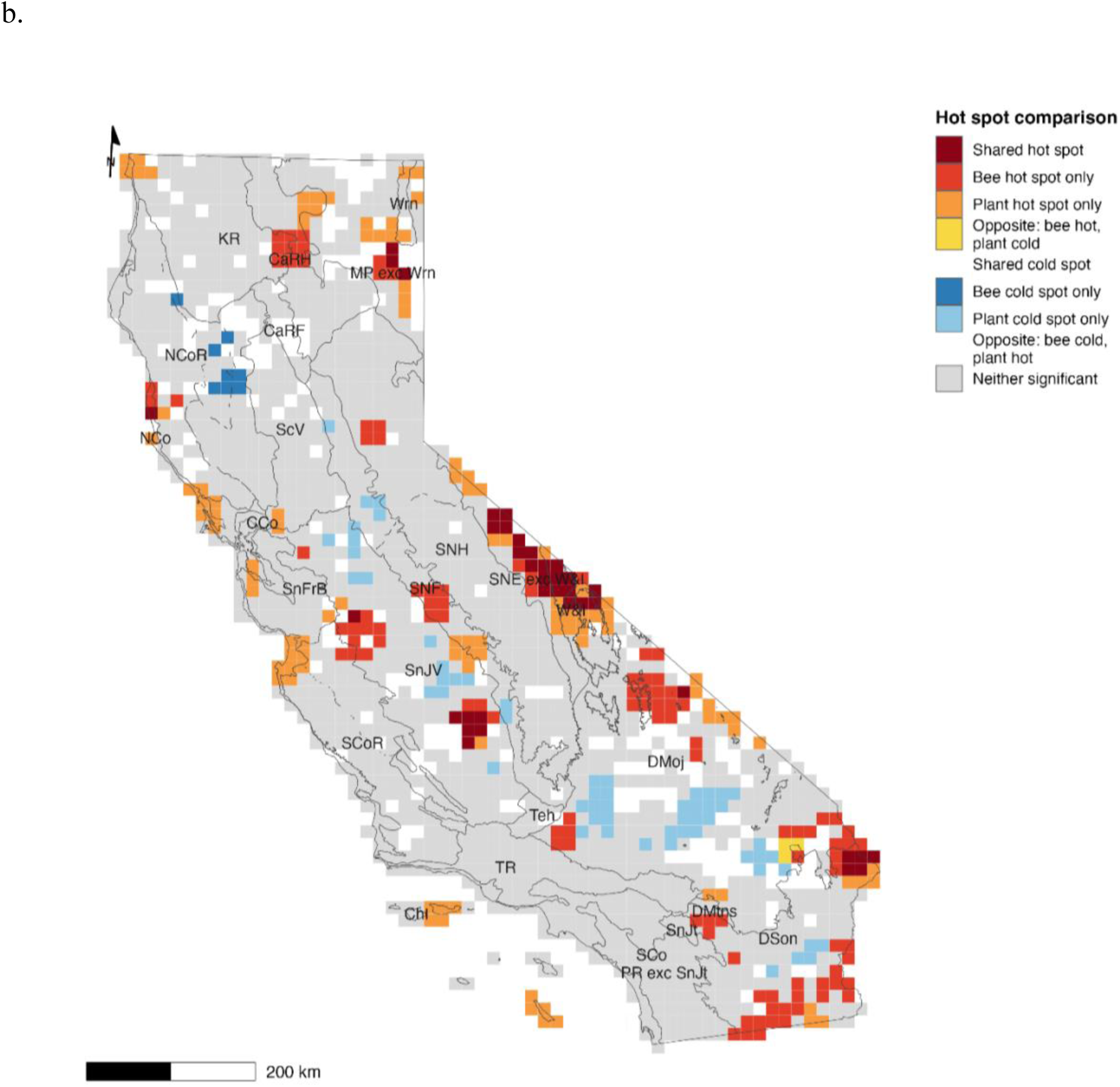
Concordance and discordance between bee and native angiosperm hot spots for (a) species richness and (b) corrected weighted endemism. Each 15 km × 15 km cell is classified by the joint outcome of the two Gi* analyses shown in Figures 7 and 8, using the same queen-contiguity bins. Angiosperm values were aligned to the bee grid before classification; cells with no matched angiosperm value are drawn in white and are not represented in the legend. Shared cold spots covered less than 1% of cells for richness and were absent altogether for corrected weighted endemism. Jepson subdivision codes are given in Table 1. Coordinate reference system: California Albers (EPSG:3310).

Angiosperm-only hotspots, where angiosperm richness was significantly high and bee richness was not significant, were found in the southern High Cascade Ranges (CaRH), northern Sierra Nevada Foothills (SnF), North Coast Ranges (NCoR), San Francisco Bay Area (SnFrB), South Coast Ranges (SCoR), the southern the northwestern Peninsular Ranges (PR), and the outermost Central Coast (CCo) and dominated much of the High Sierra Nevada (SNH), the central Klamath Ranges (KR), and the western Transverse Ranges (TR).

Bee-only hotspots, where bee richness was significantly high and angiosperm richness was not significant, occurred in scattered areas along the High Sierra Nevada (SNH) and its foothills (SNF), East of Sierra Nevada (SNE), the northern South Coast Ranges (SCoR), the eastern San Francisco Bay Area (SnFrB), the southwestern Sacramento Valley (ScV), the central Klamath Ranges (KR), Mojave Desert (DMoj), and the western Sonoran Desert (DSon) adjacent to the Peninsular Ranges. Cells in which an angiosperm coldspot coincided with a bee hotspot were the rarest outcome on the map, comprising three grid cells, all within the southern Sacramento Valley (ScV) and Mojave Desert (DMoj).

### Comparison of bee and angiosperm patterns of corrected weighted endemism

Both concordance and discordance were considerably less abundant for corrected weighted endemism (Figure 10b) than for species richness: approximately 67% of the analysis grid was not significant for either taxon, compared with 41% for richness, and shared hotspots for CWE accounted for only about 2% of cells. Areas of shared high CWE formed several discrete clusters: the largest along the area East of Sierra Nevada including the White and Inyo Mountains (SNE exc W&I, W&I); a second in the southern San Joaquin Valley (SnJV); a third on the eastern Modoc Plateau (MP exc Wrn), immediately south of the Warner Mountains; a fourth in the North Coast (NCo) and North Coast Ranges (NCoR) north of Point Arena, and a fifth in the northern Sonoran Desert near its boundary with the Mojave Desert (DSon, DMoj). Shared coldspots were absent altogether. The contrast with the richness comparison is marked: the southern California mountains that formed the dominant shared richness hotspot, comprising the Transverse Ranges (TR), the Peninsular Ranges including San Jacinto Mountains (PR exc SnJt, SnJt), and the South Coast (SCo), returned no significant CWE signal for either taxon, as did the High Sierra Nevada (SNH).

Areas of dissimilarity with high angiosperm and low bee CWE did not occur. Angiosperm-only hotspots were widely scattered, occurring East of the Sierra Nevada, including the White & Inyo Mountains (SNE exc W&I, W&I), in the eastern Mojave Desert (DMoj), eastern and southern Sonoran Desert (DSon), the Modoc Plateau (MP), Warner Mountains (Wrn), the northern Klamath and High Cascade ranges (KR, CaRH), the North and Central Coast (NCo, CCo), the northern part of the southern Sierra Nevada Foothills (SNF), the San Joaquin Valley (SnJV), and the Channel Islands (ChI).

Bee-only hotspots were strongly desert-centered, with the Mojave Desert (DMoj) and Sonoran Desert (DSon) together accounting for more than half of all such cells, and additional cells in the San Joaquin Valley (SnJV) and adjacent Sierra Nevada foothills (SNF) and northern Inner South Coast Ranges (SCoR), northern High Sierra Nevada (SNH), Cascade Ranges (CaRH), and North Coast (NCo). Cells combining an angiosperm coldspot with a bee hotspot were rare, comprising three cells, all along the eastern boundary between the Mojave and Sonoran deserts (DMoj, DSon). Angiosperm-only coldspots (approximately 3% of cells) were found in the Sierra Nevada Foothills (SNF), Mojave and Sonoran deserts (DMoj, DSon), and San Joaquin Valley (SnJV).

Simple regression analysis was used to explore the relationship between angiosperm richness and bee species richness, and between angiosperm richness and bee CWE. Angiosperm richness explained a significant portion of the variation in bee species richness (*F*(1, 1692) = 463.76, *p* < 0.0001, *R²* = 0.212], with bee richness increasing as angiosperm richness increased. Angiosperm CWE was also a statistically detectable but ecologically trivial predictor of bee CWE (*F*(1, 1692) = 13.39, *p* = 0.001, *R²* = 0.007); as the relationship explained less than 1% of variation in bee CWE.

## Discussion

While species richness and endemism of bees have been examined on a global scale^27,55^ and in regions of Greece^56^ and South Africa^57^, to our knowledge, this study is the first statewide specimen-based analysis of both bee species richness and endemism in California.

### Patterns of sampling

Bee specimen occurrence data are the best documented source of information for studying distributions even though there are biases similar to the biases in plant specimen occurrence data used here^37^. Our comprehensive analysis of 559,021 occurrence records from 162 different sources demonstrates both the breadth and limitations of available data. The dataset represents 1,742 species, exceeding recent estimates of approximately 1600 native species for California^48^, suggesting good taxonomic coverage despite acknowledged sampling gaps.

Sampling intensity varies substantially across California, with higher effort concentrated near research institutions and accessible areas. The Central Valley and southeastern deserts show sampling deficits, potentially leading to underestimates of diversity in these regions. Studies of bee species distributions using occurrence data in other states, such as Michigan, Colorado, and Texas, have been performed for conservation purposes and found significant under- representation of species in their state-level bee species list^58–61^, so the number of species found in digitized California collections likely represents a high proportion of species present in California.

Like the California plant analysis^37^, grid-cell species richness increased with sampling intensity indicating that more sampling increases the number of species detected. Achieving complete detection of all taxa in every grid cell would therefore require extremely intensive sampling and is unlikely to be an efficient use of limited conservation resources. Like collecting herbarium specimens^37^, most insect collecting is rarely equivalent to randomized plot-based sampling; occurrence datasets are often shaped by uneven effort across space and time and by targeted collecting priorities. Museum collections often concentrate on areas with high diversity and endemism, emphasize faunal or floral documentation often acquiring only a few occurrences per locality, and emphasize collecting unusual rather than common taxa^62^. These systematic biases suggest that under-sampling-driven underestimates of richness may be most pronounced in regions assumed to be species-poor, where collecting effort tends to be lowest^37^.

One of the primary challenges encountered in our study was the variability in sampling effort across different regions, which could influence the accuracy of our richness and endemism estimates. The angiosperm dataset was much larger than the completed bee dataset and had more sampled areas, with a more even distribution of sampling effort. Addressing data gaps and biases in digitized biodiversity datasets should remain a priority for bee research, as these issues can affect the accuracy and reliability of spatial analyses^30,34^. More strategic sampling and standardized monitoring methods^63^, including expanded digitization and georeferencing of collections should help to fill these data gaps^30^.

As with any gridded biodiversity analysis, results may be influenced by the choice of spatial grain; smaller cells would likely increase the number of unsampled or sparsely sampled units, whereas larger cells would smooth over local variation. While there were clearly areas that are under sampled (e.g., yellow and white cells in Figure 4 a,b), they are distributed somewhat evenly across the state of California suggesting that the reported areas of high richness and corrected weighted endemism are not misleading although there may be areas that are false negatives for high richness or false positives for low richness^37^.

### Patterns of species richness

The higher concentration of species richness for the bee fauna in the California Floristic Province and not in the Great Basin and Desert provinces is surprising given the common hypothesis that bee richness is highest in xeric areas^64–68^. The results of the present study may reflect the abundant local topographic and vegetational diversity^66^ found in the California Floristic Province. Moldenke^69,70^ found that bee diversity was lowest along the coast and in the high Sierra, moist forests of northern California, and Great Basin regions and highest in arid and semi-arid regions. Our finding of low bee species richness within the Desert Province is similarly surprising because the desert regions have been considered to be areas with high bee species richness^66,69–71^, but this could be attributed to a lack of sampling or a lack of digitization/georeferencing of collections from these areas. Our results agree with suggestions that there would be low species richness along the coast north of Los Angeles and south of the San Francisco Bay Area, the moist forests of northern California, and the northern part of the Great Basin but not some areas of the coast nearest to Los Angeles and San Francisco, the southern Great Basin (East of Sierra Nevada), or the High Sierra Nevada. Moldenke’s conclusions were based on a transect across central California and additional sites in southern California^72^ as well as specimen-based occurrence data from all of the major bee collections in California at the time of his publication^73^. Therefore, the lack of concordance with Moldenke’s data may be due to the increased availability and comprehensiveness of digital, geo-referenced specimen-based occurrence data, which provides a more complete and updated picture of bee distribution.

The weak relationship between bee and plant species richness is perhaps less surprising. Low levels of flowering plant abundance are known to have a negative effect on solitary and social bees^43,74^. Yet, only part of any flora consists of bee pollinated plants^18^ and previous studies have shown that bee and plant species richness do not always match^55,66^. The mismatch between bee and angiosperm hotspots may partly reflect differences in pollination mode, plant functional importance to bees, and variation in specialization. Restricting the plant comparison to bee-associated plant groups may be a useful direction for future study^75^.

A stronger relationship between bee and flowering plant species richness for California would be expected at a broader spatial scale of comparison, such as across North America, where much of California is a floristic hotspot^76^ and also a bee hotspot, with the high proportion of bees of North America north of Mexico found in California. A useful next step would be a continental scale analysis to provide a clear understanding of the relationship between flowering plants and bee species.

Our analysis shows that bee and plant hotspots coincide in southern California and the central coast and along the southern Sierra, while bees have independent clusters elsewhere. In addition, the Central Sierra Nevada (including Yosemite National Park), southern Sacramento Valley, San Francisco Bay Area, northern South Coast Ranges, montane Southwestern California, and the South Coast are areas of high bee biodiversity worthy of conservation attention. These areas are of ecological importance and are potentially highly vulnerable to environmental changes^77–79^.

### Patterns of endemism

Our analysis shows that bee and plant hotspots of corrected weighted endemism coincide east of the Sierra Nevada including the White and Inyo Mountains, in the southern San Joaquin Valley, on the eastern Modoc Plateau south of the Warner Mountains, in the North Coast and North Coast Ranges north of Point Arena, and in the northern Sonoran Desert near its boundary with the Mojave Desert. The areas within the California Floristic Province where bee only hotspots of bee CWE were identified were the North Coast Range, northern South Coast Ranges and adjacent San Joaquin Valley, and small regions in the southern San Joaquin Valley, central Sierra Nevada Foothills, northern High Sierra Nevada, and eastern Klamath and adjacent Cascade Ranges. Numerous other hotspots of corrected weighted endemism were found within the Great Basin and Desert provinces, including the Modoc Plateau, the eastern Sierra Nevada, northern and eastern Mojave Desert, the Desert mountains, and southern Sonoran Desert along borders with Arizona and Mexico. The Warner Mountains subregion was the only subregion of transmontane California without such a hotspot. These bee-only regions represent areas of unique biodiversity in California that would be missed by analyses based on plant data alone.

*Spatial structure of diversity patterns*.

The contrasting spatial patterns between species richness and endemism reveal fundamental differences in how these diversity metrics are distributed across California’s landscape. The moderate spatial autocorrelation in species richness (Moran’s *I* = 0.300) reflects broad environmental gradients (e.g., temperature, rainfall, vegetation productivity) that create large, coherent regions of high and low diversity. These gradients operate at regional to landscape scales and may produce the smooth spatial structure we observed. In contrast, the weak spatial autocorrelation in corrected weighted endemism (Moran’s *I* = 0.089) does not indicate the absence of spatial structure, but rather a different kind of structure: endemism is concentrated in a relatively small number of high-value cells embedded within a landscape that is otherwise low in endemism relative to overall richness. This pattern is consistent with ecological theory: high endemism often occurs in isolated areas with unique combinations of microhabitat, geology, or climatic conditions that do not necessarily correlate with the factors driving overall species richness^80,81^.

The Moran scatterplots further illustrate this distinction (Figure 9). The extreme compression of CWE values near zero for most grid cells, combined with a small number of cells with standardized endemism values exceeding 10–25X, is the visual signature of localized endemism hotspots. The pattern of extreme right-skew in the CWE distribution confirms that a handful of grid cells harbor disproportionately high endemism relative to their neighbors. These outliers may represent areas with specialized conditions—perhaps unique microclimates or historical refugia—that support distinct endemic assemblages while being surrounded by cells with low endemism relative to richness. This fine-scale spatial heterogeneity in endemism has important conservation implications: protecting biodiversity hotspots based solely on species richness may miss critical areas harboring concentrations of rare, range-restricted taxa. Our findings suggest that effective conservation strategies must account for both broad-scale richness gradients and fine-scale endemism hotspots, as these patterns operate at different spatial scales and likely respond to different environmental drivers.

### Regional patterns Southwestern California

The hotspot analysis performed for angiosperm species richness aligns with the detected high native vascular plant species richness throughout the montane areas of Southwestern California and the South Coast. Baldwin et al.^37^ identified the Northern and Southern Channel Islands as hotspots of plant species richness but in our gridded Getis-Ord *Gi\** analysis this pattern is weaker. Because bees are attracted to plants that produce pollen and nectar, we expected that many areas of high plant richness would also support high bee species richness. Our findings show broad correspondence between plant and bee richness in hotspots in parts of Southwestern California, but this correspondence is incomplete. Bee richness hotspots are not as evident in the Western Transverse Ranges and Channel Islands. In contrast, while Baldwin et al.^37^ found areas of high angiosperm corrected weighted endemism on the Channel Islands and within the San Bernardino Mountains, bee CWE hotspots were not broadly developed in coastal Southwestern California or on the Channel Islands, although localized inland or desert-edge hotspots occur nearby.

### Central Western California

The finding of large hotspots for bee species richness within Central Western California is not surprising due to the long history of bee research and the density of research universities within this region^60,72,82–85^. Despite these studies, bee corrected weighted endemism hotspots are much more limited than bee richness hotspots, with the only cluster occurring within the Inner South Coast Ranges. Bee and plant richness hotspots overlap in parts of Central Western California, especially in the San Francisco Bay Area, but bee and plant richness hotspots are not spatially identical. The diversity of flowering plants outside angiosperm hotspots may still be quite high, though, especially by comparison to areas outside California. It must be remembered that the comparisons here are within the context of Californian plant diversity. If examined on a national scale, much of California would be viewed as a hotspot for plant diversity (see Figure 2 in Mishler et al.^76^). As with bees, the San Francisco Bay Area and Central Coast have strong angiosperm richness hotspots but few corresponding angiosperm corrected weighted endemism hotspots, similar to results for Southwestern California, where unusually high richness, rather than a low number of narrowly restricted taxa, results in low CWE.

### Northwest California

Bee species richness hotspots were rare in Northwest California, limited to two isolated cells in the Klamath Ranges, despite the presence of hotspots for angiosperm species richness in the Klamath Ranges and Inner North Coast Ranges. This difference between bee and plant diversity in Northwest California is not too surprising as previous studies found this region had low bee species richness^86^ while both overall and serpentine endemic Californian plant diversity increased with increases in latitude and mean annual rainfall^87^. Bee and plant corrected weighted endemism hotspots are scarce in Northwest California, and corrected weighted endemism hotspots are more localized, occurring mainly in far northern or immediate coastal areas. This is the wettest part of California. Previous studies have suggested relationships between precipitation patterns and bee diversity, with bee richness potentially higher in more xeric conditions^66–68^, while plant diversity often responds positively to precipitation^66,87^.

However, the specific climatic drivers of diversity patterns in California warrant detailed future investigation using comprehensive environmental datasets, as mentioned earlier in the Discussion.

### Cascade Ranges

No hotspots for bee species richness were found within the Cascade Ranges. A small hotspot for bee corrected weighted endemism could be found within the High Cascade and adjacent Klamath ranges. As in Northwest California, hotspots for angiosperm species richness were found in the Cascades. The eastern Cascades and western Modoc Plateau were notable coldspots for angiosperm species richness. Angiosperm corrected weighted endemism hotspots are rare, with small hotspots occurring near the far northern High Cascade Ranges.

### Sierra Nevada

Bee species richness hotspots occur patchily within the Central High Sierra Nevada and parts of the Central Sierra Nevada Foothills with smaller areas of high bee species richness within the Northern High Sierra Nevada, Northern Sierra Nevada Foothills, and Southern High Sierra Nevada. This is another case where the high level of research on solitary and social bees in this region^88–90^ may bias the results. No large hotspots were found for bee corrected weighted endemism within the Sierra Nevada proper; but hotspots are found east of the range. It is surprising that bee species richness hotspots are less extensive than angiosperm species richness through much of the High Sierra Nevada. These findings suggest that other factors, such as soil type, may limit the positive response of pollinators to floral species richness hypothesized for this area^91^.

### Great Valley

Together, the San Joaquin Valley and Sacramento Valley make up the Great Valley, also known as the Central Valley. The relatively xeric San Joaquin Valley has localized hotspots of high bee endemism but low species richness, although the northwestern part of the valley falls within a bee richness hotspot. In contrast, the relatively mesic Sacramento Valley has low endemism but contains several bee richness hotspot cells. The Great Valley has gone through many significant changes over the years including the conversion of wetlands, grasslands, and woodlands to agriculture, loss of groundwater^92^, changes in soil salinity due to climate change^93^, and increased pesticide use^94^. Groundwater is an important resource for bees and pollinator-reliant plants^95^, soil salinity can impact plant survival and can influence the attractiveness of plants to insect pollinators^96^, and the use of pesticides can have negative health effects on bees^97^. The identification of coldspots in California’s highly productive agricultural regions, particularly the San Joaquin Valley, is critical for understanding pollination vulnerabilities in one of the nation’s most important food-producing areas^98^. These gaps in pollinator populations could significantly impact crop yields and agricultural sustainability, making it essential to investigate the underlying causes and develop targeted conservation strategies to support both wild and managed bee populations in these intensively farmed landscapes.

### Modoc Plateau

No hotspots for bee species richness were found within the Modoc Plateau. Small and localized hotspots for bee corrected weighted endemism were found there. There were also no broad hotspots for angiosperm species richness. In contrast, hotspots for angiosperm corrected weighted endemism were extensive within the Modoc Plateau relative to the hotspots for bee corrected weighted endemism. It is not surprising to find a lack of corresponding bee and flowering plant species richness hotspots due to the ecosystems within the Modoc Plateau being mostly semi-arid sagebrush rangelands, shrub steppes, and juniper woodlands^99^.

### East of Sierra Nevada

Hotspots for bee species richness within the East of Sierra Nevada region were present but limited in extent. Hotspots for bee corrected weighted endemism were widespread but patchy across the region, including the White and Inyo Mountains and had significant concordance with angiosperm CWE hotspots. Baldwin et al.^37^ explored whether this pattern of plant endemism was influenced by species that are more widely distributed in Nevada than in California by restricting the analyses to absolute California endemics that do not extend outside California. They found that the East of the Sierra Nevada pattern of high corrected weighted endemism still held, but the pattern did not hold in the Modoc Plateau, where it does appear that range-restricted taxa within California that extend in distribution outside of the state are driving the CWE pattern.

### Deserts of California

Bee species-richness hotspots were limited in the Mojave Desert, and angiosperm species-richness hotspots were absent or weak there. The inclusion of the California Floristic Province in these analyses places diversity patterns in the cold Great Basin and warm Mojave and Sonoran deserts of California in direct comparison with a global-scale biodiversity hotspot^100^. This statewide framing is important in discerning whether deserts appear depauperate because they support fewer taxa, or because their richness is being evaluated alongside one of the most species-rich and endemic regions in North America. By comparison with deserts elsewhere in southwestern North America, substantial parts of the California deserts appear quite rich and high in plant endemism^76^. Hotspots for bee and angiosperm corrected weighted endemism were present, but their spatial patterns differed. Bee corrected weighted endemism hotspots were relatively prominent and patchily distributed in the Mojave and Sonoran Desert regions, whereas angiosperm corrected weighted endemism hotspots were more restricted. A modest number of hotspots for bee species richness were found within the western Sonoran Desert, but most of these hotspots were shared with Southwestern California. In contrast, there were no broad hotspots for angiosperm species richness in the eastern Mojave and eastern Sonoran Desert regions. The far eastern Mojave Desert has extensive hotspots for angiosperm corrected weighted endemism. The concentration of specialist bee species may explain why bee species richness hotspots occur in these areas where angiosperm richness hotspots do not, as specialists can maintain populations even during periods of reduced floral diversity^101^. In addition, bee detectability and abundance in arid regions may vary strongly among years, which may make desert diversity more difficult to characterize from historical specimen data.

### Comparison of bees and angiosperms

Bee and angiosperm diversity patterns exhibited both similarities and dissimilarities. While the observed associations of richness between bees and flowering plants do not support the notion that bee distributions are tightly coupled to the diversity of floral resources^42^, the regression analysis results suggest that plant species richness does play a role in supporting diverse bee communities. Angiosperm richness was also a statistically significant predictor of bee corrected weighted endemism, however, the relationship explained very little variation in bee corrected weighted endemism. Together, these results suggest that, although bees and plants may share some broad distributional signals, additional factors shape their spatial patterns. Variation in specialization among bees and flowering plants, and differences among plant species in their importance as nectar and pollen resources, likely weaken expected relationships between angiosperm and bee richness and endemism^18,72^. At broader (e.g., continental) spatial scales, much of California stands out as exceptionally rich and endemic floristically, but comparable large-scale spatial evaluations have not yet been conducted for bees. Going forward, understanding the effect of climate change on plant distributions may be more significant to bee distributions — and pollinator conservation more generally — than understanding its impact on plant biodiversity *per se*^102^. It is also essential to acknowledge that multiple factors, such as climate, habitat complexity, and nesting resources, can independently influence bee distribution patterns^41^. In addition, bee host-plant specialization is a plausible mechanism underlying the discordance between bees and plants and may be an important direction for future work. Future studies should explore the effects of these additional variables to gain a more comprehensive understanding of bee biodiversity.

The modest correlation between bee and plant species richness (*R²* = 0.21) likely reflects the complex relationships between flowering plant diversity and bee community structure. Not all angiosperms rely on bee pollination; data suggests that approximately 30% of angiosperm species utilize wind pollination, self-pollination, or specialized non-bee pollinators^18^. Additionally, plant species vary greatly in their value as pollen and nectar sources, with some providing critical resources for specialist bees while others may be rarely visited. The degree of plant-pollinator network specialization varies across California’s diverse ecosystems^72^, potentially explaining regional differences in bee-plant concordance patterns.

The use of hotspot analysis for conservation decision-making must be done judiciously, especially for less studied organisms as it is unknown how well hotspots identified based on data from one group of organisms can be extended to others^103,104^. Critics warn that decision makers should not think of hotspot conservation as the sole remedy for biodiversity loss^105^. Still, this information is important for identifying sites that require strong conservation efforts to protect extensive biodiversity^106^.

Regions where bee and plant hotspots coincide are the most efficient targets for conservation investment. The East of the Sierra Nevada and the White and Inyo Mountains stand out as hotspots for angiosperm and bee richness *and* for endemism in both taxa and should rank among the state’s highest priorities. The South Coast and San Jacinto Mountains share high richness, while the Peninsular Ranges and Central Coast are disproportionately important for endemism in both groups. Because these areas concentrate the diversity of both partners in the pollination mutualism, protecting them returns benefits that single-taxon planning would miss.

This study offers insight into the relationship between bee and plant biodiversity. By examining patterns of species richness and endemism for California bees, we can assess the extent to which bee populations are adequately protected by plant conservation initiatives. Our findings suggest that bees will not be well protected by conservation plans developed exclusively with angiosperm data, especially in regions of high bee endemism. Endemic bee species are often at higher risk of extinction due to specialized habitat requirements^107^ and warrant independent conservation attention. Our analysis highlights the importance of expanding the data used in conservation efforts to include not only plant diversity but also the available bee data.

Furthermore, this study underscores the need for increased attention to significant under-protected areas in the California Floristic, Great Basin, and Desert provinces in conservation planning^108^. The high endemism and species richness found in these areas highlight the importance of protecting these habitats for maintaining biodiversity^109^. By broadening the organismal scope of conservation efforts, we can better safeguard the rich biodiversity of California.

Limitations of the study

Several limitations must be acknowledged in interpreting our results. First, museum collection data contain inherent biases related to collector preferences, accessibility, and institutional priorities that may not reflect true abundance or distribution patterns^30,34^. Our sampling adequacy analysis reveals gaps in geographic coverage, particularly in remote areas and private lands, potentially leading to underestimates of diversity in under-sampled regions^30,34^. The redundancy index identifies where sampling is adequate to support the patterns we report, but they cannot recover species from cells that were never sampled, and no post hoc weighting or spatial thinning was applied to compensate for uneven effort. These biases are shared, at least in part, with the California plant occurrence data against which we compare the bee patterns^37^, so concordance between the two groups may partly reflect a common collecting history rather than shared ecology alone.

Second, assembling occurrence data from three databases that differ in field structure, taxonomic treatment, and cataloguing convention imposes its own constraints. Records lacking a catalogued identifier were excluded regardless of the quality of their locality or taxonomic data, so collections that have not assigned or exposed catalog numbers are under-represented. Duplicate detection depends on catalog numbers being recorded consistently across databases; where they are not, a specimen held in two sources may be retained twice.

Third, the 15 km grid resolution, while appropriate for landscape-level analysis and essential for comparison with Baldwin et al.^37^, may obscure fine-scale diversity patterns and microhabitat preferences important for some bee species. Fourth, our analysis does not account for temporal changes in bee communities, seasonal phenology, or recent population declines that may affect contemporary diversity patterns. Finally, taxonomic uncertainties and ongoing systematic revisions may affect species-level identifications; because verbatim determinations were harmonized against a global bee taxonomy, with synonyms replaced by accepted names, the species total we report reflects that harmonized treatment rather than the determinations on specimen labels.

Future directions

Future research priorities include detailed climatic analysis to understand environmental drivers of the diversity patterns identified here, examination of temporal changes in bee communities, and investigation of fine-scale habitat relationships that may explain local diversity hotspots.

A spatial phylogenetic framework would extend the analysis in a further direction^76,110^. Species richness and corrected weighted endemism treat all species as equivalent units, capturing the number and range-restriction of taxa but not the evolutionary distinctiveness of the lineages they represent, so two cells with identical richness may differ substantially in the amount of phylogenetic history they hold. Incorporating evolutionary history would deepen our understanding of the conservation value of different regions^110^.

## Resource availability

**Data and code:** Requests for further information and resources should be directed to and will be fulfilled by the lead contact, Gretchen Le Buhn. All scripts and the analytical dataset of California bee occurrence records (ca_bees.csv), the Biodiverse output shapefiles (ca_bees_shapefile.shp, ca_plants_shapefile.shp), the archived global bee taxonomy reference (beesTaxonomy_2026-05-07.Rda), and the complete institutional attribution, taxonomic identifier, collector, and species lists have been deposited at Zenodo (https://doi.org/10.5281/zenodo.15653300) and are publicly available as of the date of publication. The DOI is listed in the key resources table. Occurrence records from the USDA-ARS Pollinating Insects Research Unit are not held in a public repository and are available on request from that unit; the contact is listed in the key resources table. The plant data are available from Dryad (https://doi.org/10.6078/D16K5W ) or the UC Berkeley Dash repository (https://dx.doi.org/10.6078/D1G010 ), as detailed by Baldwin et al.^37^. Grid generation and diversity metric calculation were performed interactively in Biodiverse v4.3 and are therefore not scripted; the Biodiverse output shapefiles are deposited so that all downstream analyses can be reproduced without re-running Biodiverse.

## Materials availability

This study did not generate new unique reagents or materials.

## Acknowledgments

We would like to thank Jerry Davis for advice and suggestions on the hotspot analysis and for his feedback on the thesis. We thank Harold Ikerd for sharing data from the USDA-ARS Pollinating Insects Research Unit. We particularly acknowledge the Big-Bee project team and the 13 collaborating institutions that have made bee digitization data widely available for research. We thank the U.S. National Park Service and the following park units for providing access to specimen data from their museum collections: Channel Islands National Park (Accession CHIS-536); Death Valley National Park (Accessions DEVA-2649 and DEVA-2616); John Muir National Historic Site (Accession JOMU-493); Joshua Tree National Park (Accession JOTR-1006); Mojave National Preserve (Accession MOJA-182); Pinnacles National Park (Accessions PINN-458, PINN-480, PINN-482, PINN-519); Point Reyes National Seashore (Accession PORE-885); Redwood National Park (Accession REDW-325); Santa Monica Mountains National Recreation Area (Accession SAMO-293); and Yosemite National Park (Accession YOSE-6887).We gratefully acknowledge the 162 institutions and countless individuals who contributed specimen-based occurrence data to this analysis (Table S4, complete list in Zenodo). We thank the many collectors, taxonomic specialists, and database managers whose efforts made this analysis possible (Tables S5 and S6, complete lists in Zenodo).

## Author contributions

GA, GL, and BB conceived the concept of the study. GA and GL collected and analyzed the data. GL and SJ directed the project. KS provided informatics expertise, data validation, and insights from the Big-Bee digitization project. GA, GL, BB, and SJ prepared the manuscript. All authors contributed to manuscript revision and approved the final version.

## Supplemental Information

Document S1. Figures S1–S2 and Tables S1–S7

**Figure S1.**
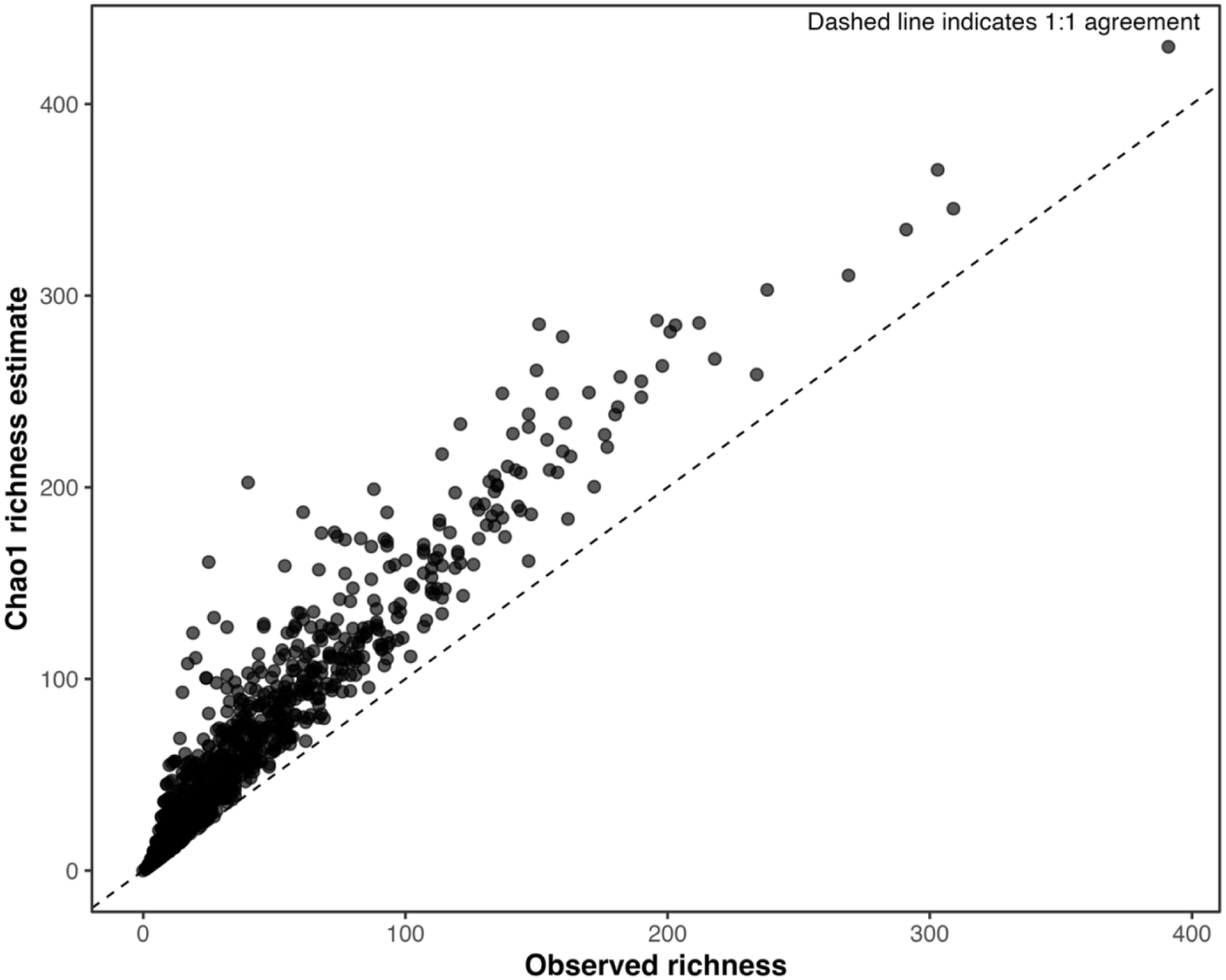
Observed species richness compared with Chao1-estimated richness across California bee grid cells, related to Figures 5a, 6a and 7a. Each point represents one 15 km × 15 km grid cell (n = 1,694). The dashed line indicates 1:1 agreement between observed richness and Chao1- estimated richness. Points above the line indicate cells in which estimated richness exceeded observed richness, consistent with incomplete sampling. The two metrics track one another strongly across the richness gradient, although Chao1 generally produces higher estimates.

**Figure S2.**
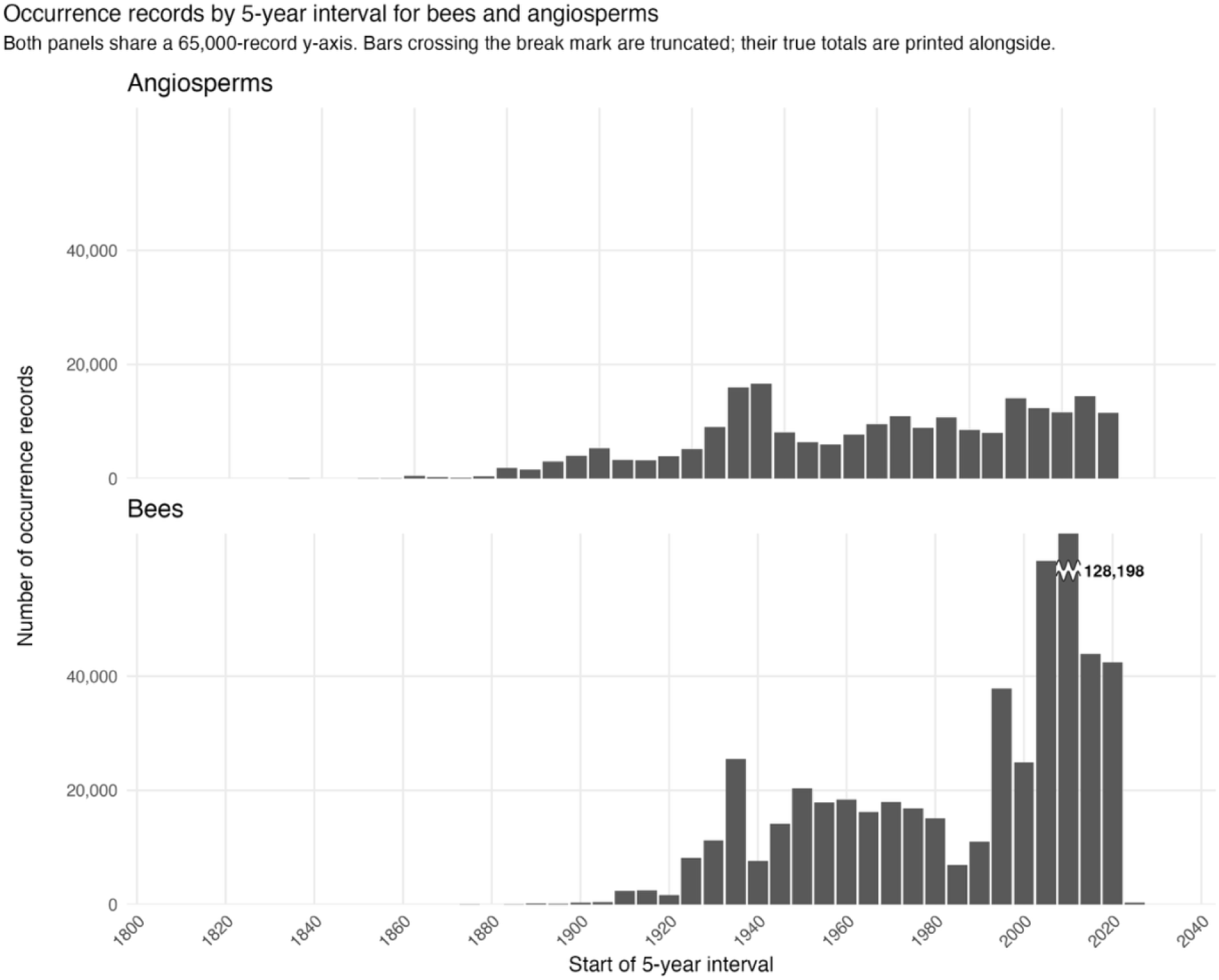
Occurrence records per five-year interval for California angiosperms and bees, related to STAR Methods. Bars show the number of records per five-year collection interval for the cleaned bee dataset (lower panel) and the angiosperm dataset (upper panel), from 1800 onward; intervals are labelled by their starting year. Both panels share a common y-axis ceiling so that sampling effort is directly comparable between taxa. Bars exceeding that ceiling are truncated at a break mark, with the true interval total printed alongside in bold. Two bee records dated 1700 fall outside the plotted range.

**Table S1.**
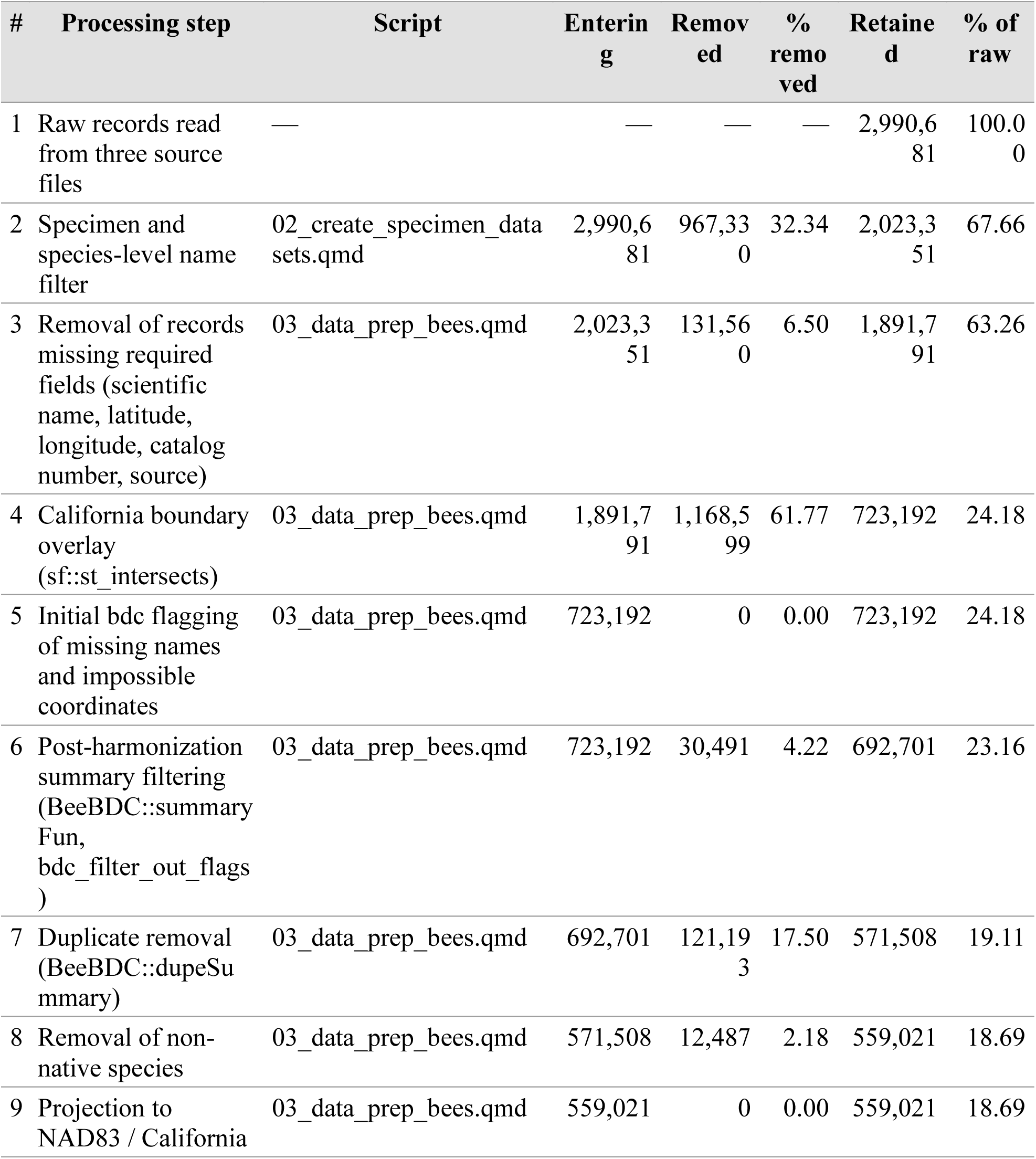
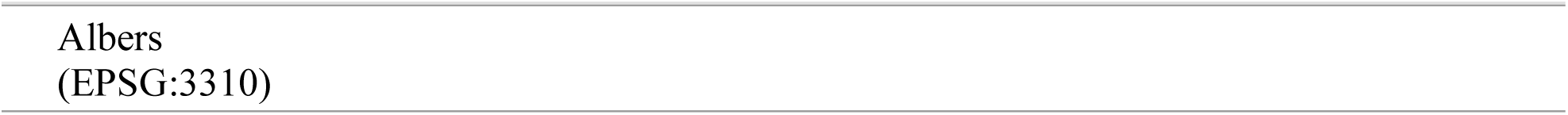
Record attrition from raw source files to the final analytical dataset, related to STAR Methods. Each step operates on the output of the step above it. Raw records by source: USDA/BBSL 120,516; Big Bee 510,033; Dorey 2,360,132. Step 5 removed no records because the completeness filter at step 3 had already excluded records lacking names or coordinates. The final analytical dataset contains 559,021 occurrence records representing 1,742 species, or 18.69% of the raw records read from the three source files.

**Table S2.** Records contributed and retained by each source database, related to STAR Methods. Counts are taken at the point of de-duplication and compared with records surviving into the final analytical dataset. The retention rate reflects the sourceOrder priority rule and should not be read as a measure of data quality. Sources are the Big Bee project [S1], the globally synthesised bee occurrence dataset of Dorey et al. [S2], and the USDA-ARS Pollinating Insects Research Unit.

| **Source** | **Records contributed** | **% contributed** | **Records retained** | **% of retained** | **Retention rate (%)** |
| --- | --- | --- | --- | --- | --- |
| Big Bee | 338,294 | 48.8 | 220,548 | 39.5 | 65.2 |
| Dorey | 235,181 | 34.0 | 221,475 | 39.6 | 94.2 |
| USDA/BBSL | 119,226 | 17.2 | 116,998 | 20.9 | 98.1 |

**Table S3.**
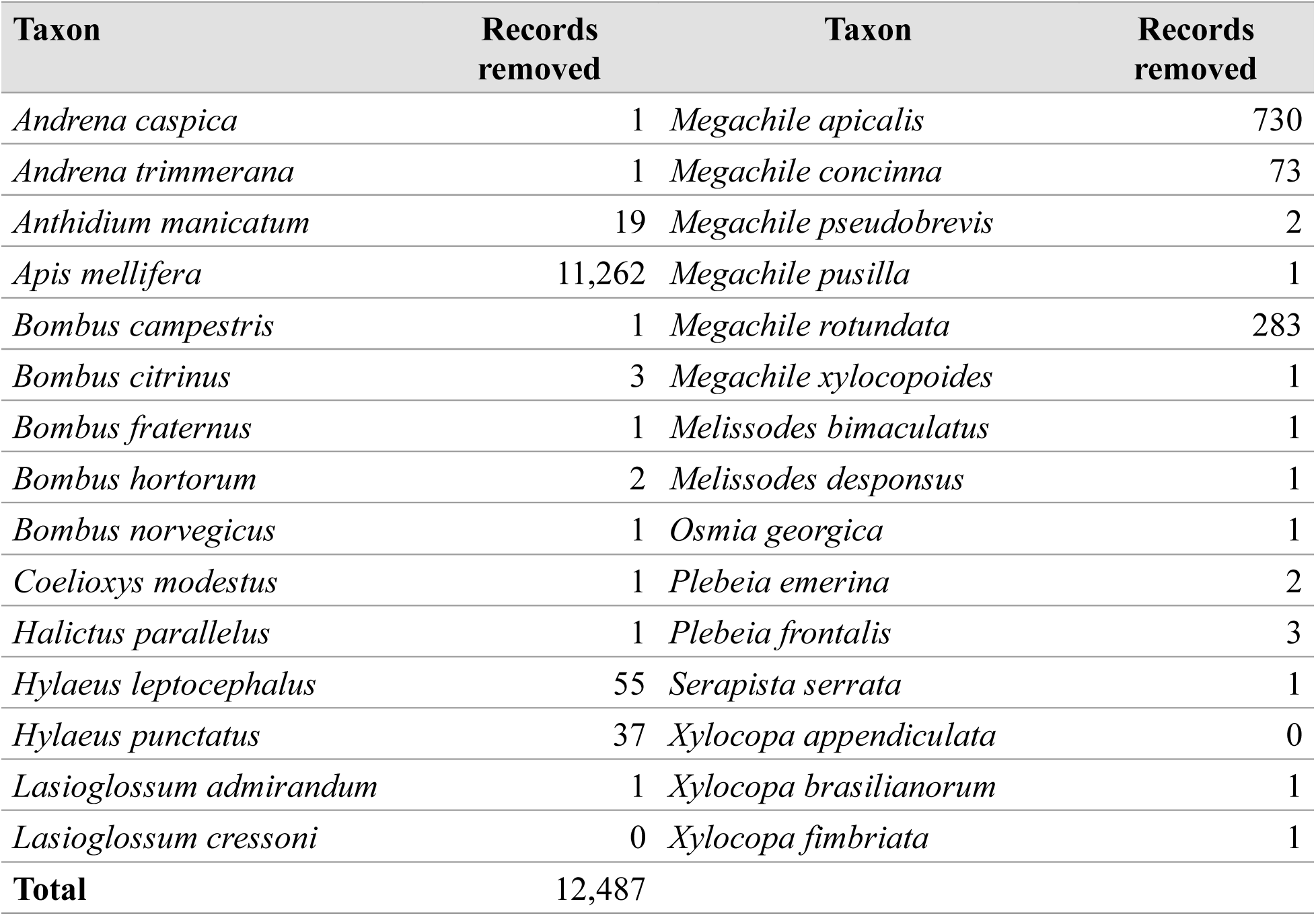
Non-native bee taxa excluded from the analytical dataset, related to STAR Methods. Records of bee species not native to California were removed after de-duplication and before coordinate projection, so that the projected analytical dataset represents the native California bee fauna. Removal was by exact match against the taxon names listed below. This step removed 12,487 records (2.2% of the de-duplicated dataset), retaining 559,021. Two listed taxa, *Lasioglossum cressoni* and *Xylocopa appendiculata*, matched no records in the cleaned dataset.

**Table S4.**
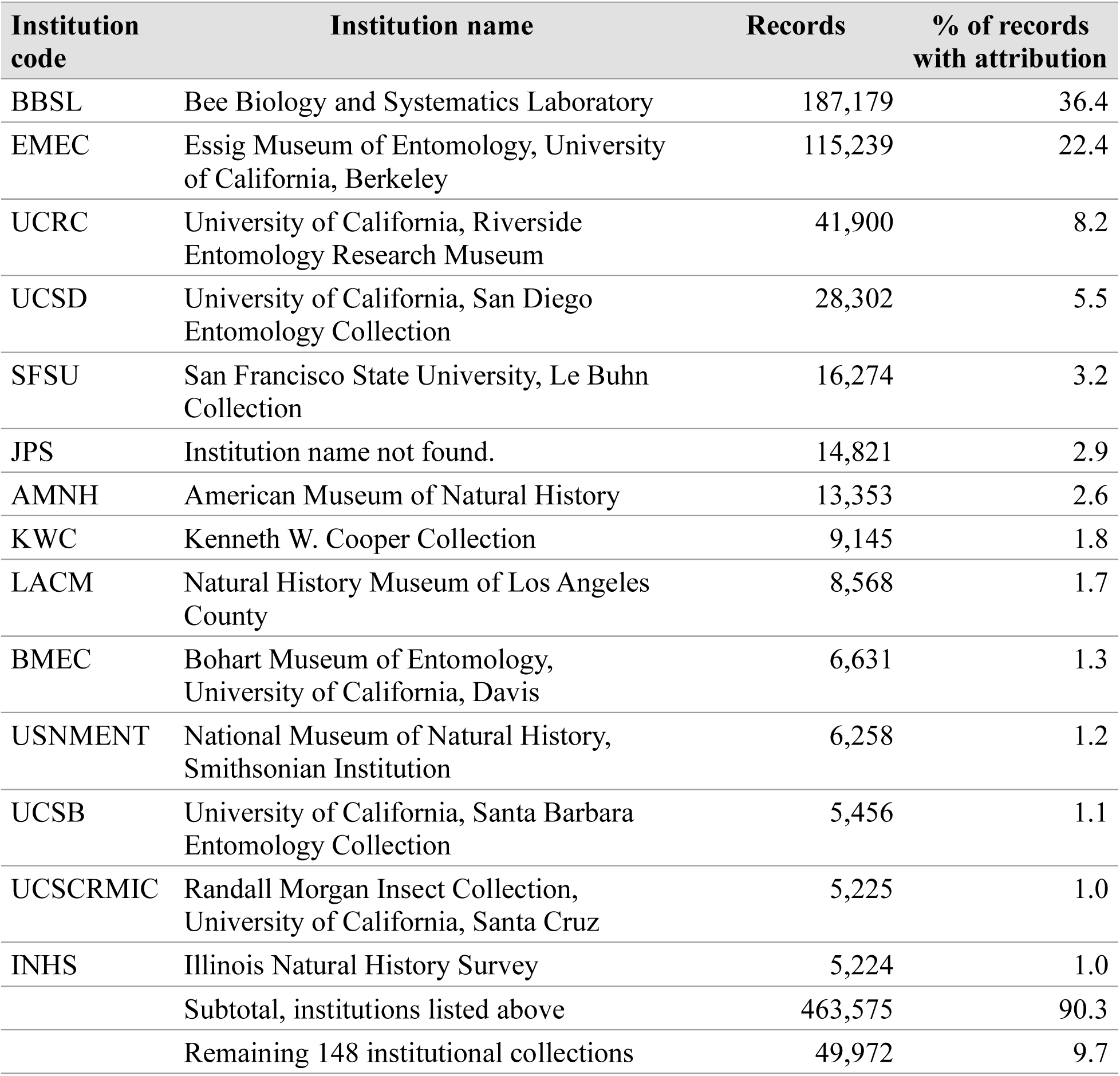
Institutions contributing at least 1% of specimen-based bee occurrence records with institutional attribution, related to STAR Methods. Institutional codes could be parsed for 513,547 of the 559,021 records in the analytical dataset (91.9%), representing 162 distinct institutional sources. Percentages are calculated against the 513,547 records carrying institutional attribution. Bee Biology and Systematics Laboratory records are aggregated across the BBSL, BBSLID, PINN, and YOSE collection and project codes. The remaining 148 institutional collections together contributed 49,972 records (9.7%); the complete list of all 162 institutions is deposited with the analytical dataset and analysis code (see data and code availability).

**Table S5.** Taxonomic identifiers with the largest numbers of identifications, related to STAR Methods. Not all occurrence records carried an identifier name. Approximately 400 unique identifier names appear in the dataset. The complete list is deposited with the analytical dataset and analysis code (see data and code availability). Name strings were not disambiguated across spelling and abbreviation variants, so counts for individuals whose names appear in more than one form are underestimates.

| Rank | Taxonomic identifier | Identifications |
| --- | --- | --- |
| 1 | T. Griswold | 76,934 |
| 2 | S. Burrows | 25,335 |
| 3 | Keng-Lou James Hung | 17,937 |
| 4 | R. Thorp | 15,275 |
| 5 | E. Frehner | 12,129 |
| 6 | H. Ikerd | 10,219 |
| 7 | J. Mullins | 8,609 |
| 8 | K. Huntzinger | 6,693 |
| 9 | B. Bagot | 6,148 |
| 10 | J. Gibbs | 5,648 |
| 11 | J. Koch | 4,814 |
| 12 | C. Stragar | 4,307 |
| 13 | A. Lehner | 4,002 |
| 14 | M. C. Orr | 3,860 |
| 15 | E. Stephens | 3,832 |
| 16 | J. Pawelek | 3,714 |
| 17 | L. Ikerd | 3,698 |
| 18 | M. Rightmyer | 2,984 |
| 19 | N. Entry | 2,795 |

**Table S6.** Collectors and collecting teams with the largest numbers of specimens, related to STAR Methods. Not all occurrence records carried a collector name. Labels bearing more than one collector were treated as a collecting team and were not added to individual collector counts, so these values are underestimates for some collectors. Name strings were not disambiguated across spelling and abbreviation variants; *J. M. Meiners and J. Meiners are probable variants of the same name and have not been merged. They would be ranked 1 if this occurred. The dataset contains 4,857 unique collector names or collecting teams; the complete list is deposited with the analytical dataset and analysis code (see data and code availability).

| Rank | Collector or collecting team | Specimens |
| --- | --- | --- |
| 1 | T. Griswold | 26,593 |
| 2 | J. M. Meiners* | 23,442 |
| 3 | Keng-Lou James Hung | 17,648 |
| 4 | O. Messinger | 15,136 |
| 5 | A. M. E. Lehner | 15,121 |
| 6 | J. Meiners* | 11,930 |
| 7 | K. Cutler | 11,718 |
| 8 | H. Ikerd | 11,690 |
| 9 | P. Timberlake | 9,189 |
| 10 | C. Michener | 9,182 |
| 11 | J. Mullins | 9,076 |
| 12 | K. Cooper | 8,971 |
| 13 | S. Kaiser | 5,688 |
| 14 | R. Hanifin | 5,591 |
| 15 | R. Morgan | 5,458 |
| 16 | E. Stephens | 5,033 |
| 17 | B. H. Bagot | 5,008 |
| 18 | R. Jaffe | 4,889 |
| 19 | K. Ullmann | 4,584 |
| 20 | L. Macior | 4,470 |

**Table S7.**
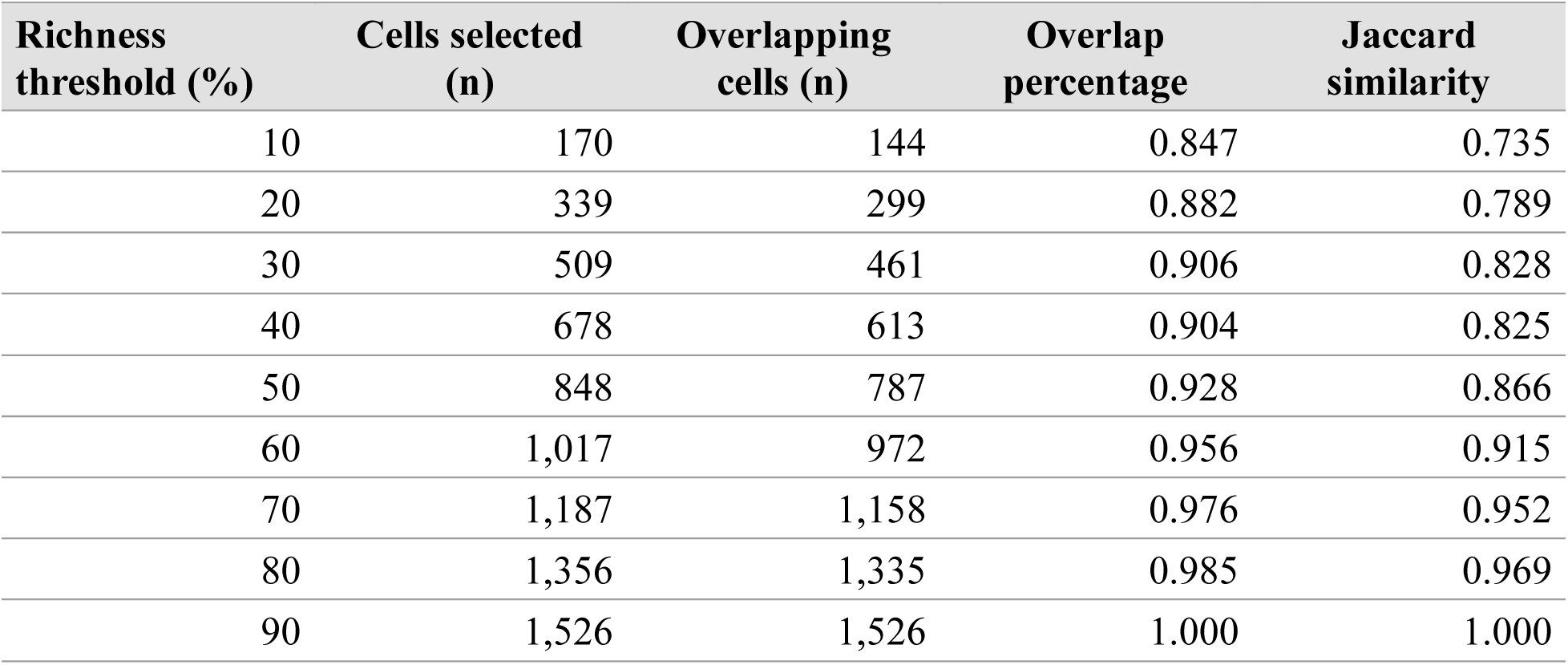
Overlap between grid cells ranked by observed species richness and by Chao1- estimated richness related to Figure 5a, 6a and 7a. Threshold percentages indicate the proportion of all 1,704 grid cells retained after ranking cells separately under each metric. Cells selected is the number of cells included in each top-ranked set, overlapping cells is the number shared between the two sets, and overlap proportion is the fraction of selected cells shared between them. High overlap indicates that observed richness and Chao1 identify broadly similar sets of relatively high-value cells.

| **Richness threshold (%)** | **Cells selected (n)** | **Overlapping cells (n)** | **Overlap percentage** | **Jaccard similarity** |
| --- | --- | --- | --- | --- |
| 10 | 170 | 144 | 0.847 | 0.735 |
| 20 | 339 | 299 | 0.882 | 0.789 |
| 30 | 509 | 461 | 0.906 | 0.828 |
| 40 | 678 | 613 | 0.904 | 0.825 |
| 50 | 848 | 787 | 0.928 | 0.866 |
| 60 | 1,017 | 972 | 0.956 | 0.915 |
| 70 | 1,187 | 1,158 | 0.976 | 0.952 |
| 80 | 1,356 | 1,335 | 0.985 | 0.969 |
| 90 | 1,526 | 1,526 | 1.000 | 1.000 |

## STAR★METHODS

*Patterns of species richness and endemism in bee and plant communities in California*

### KEY RESOURCES TABLE

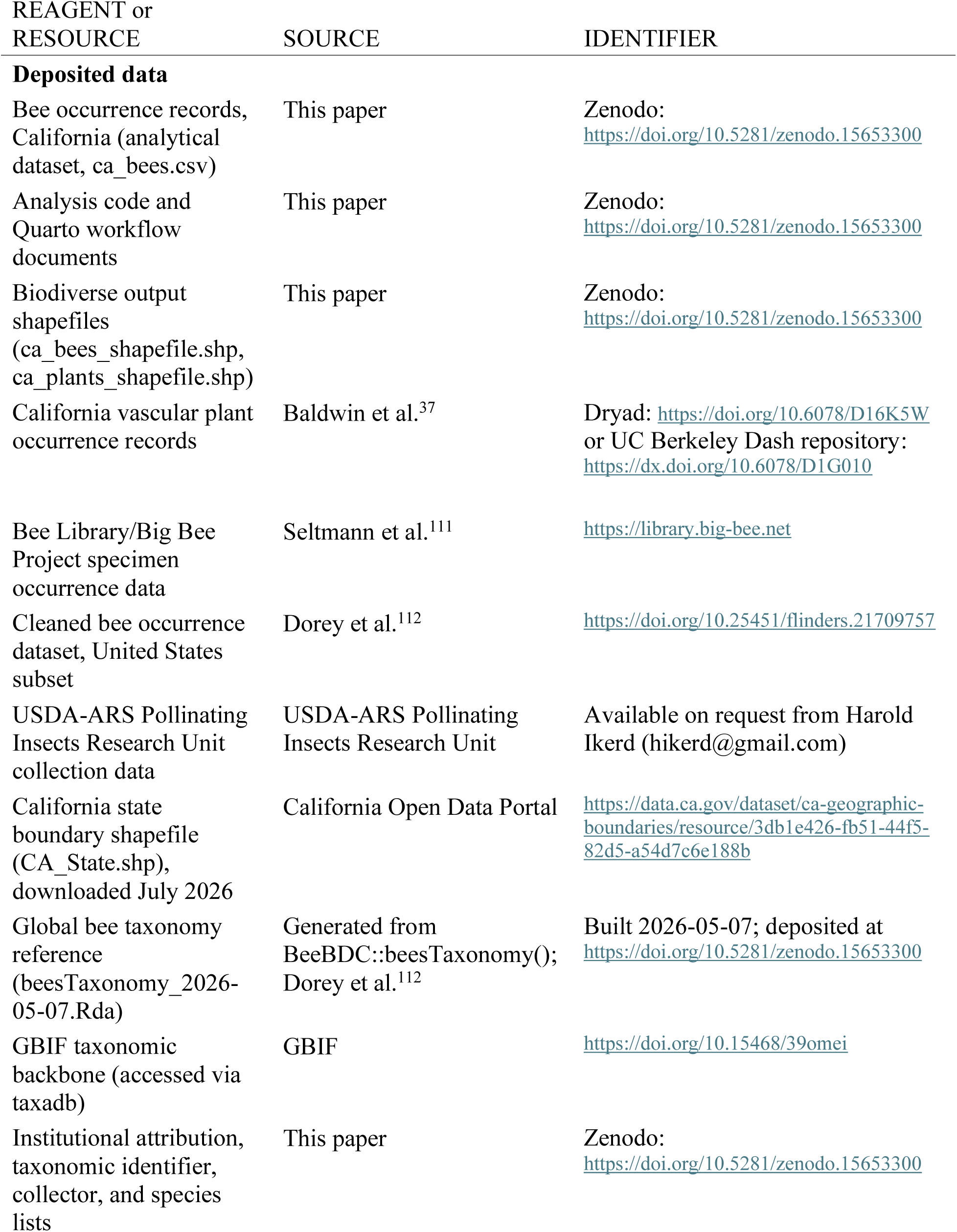
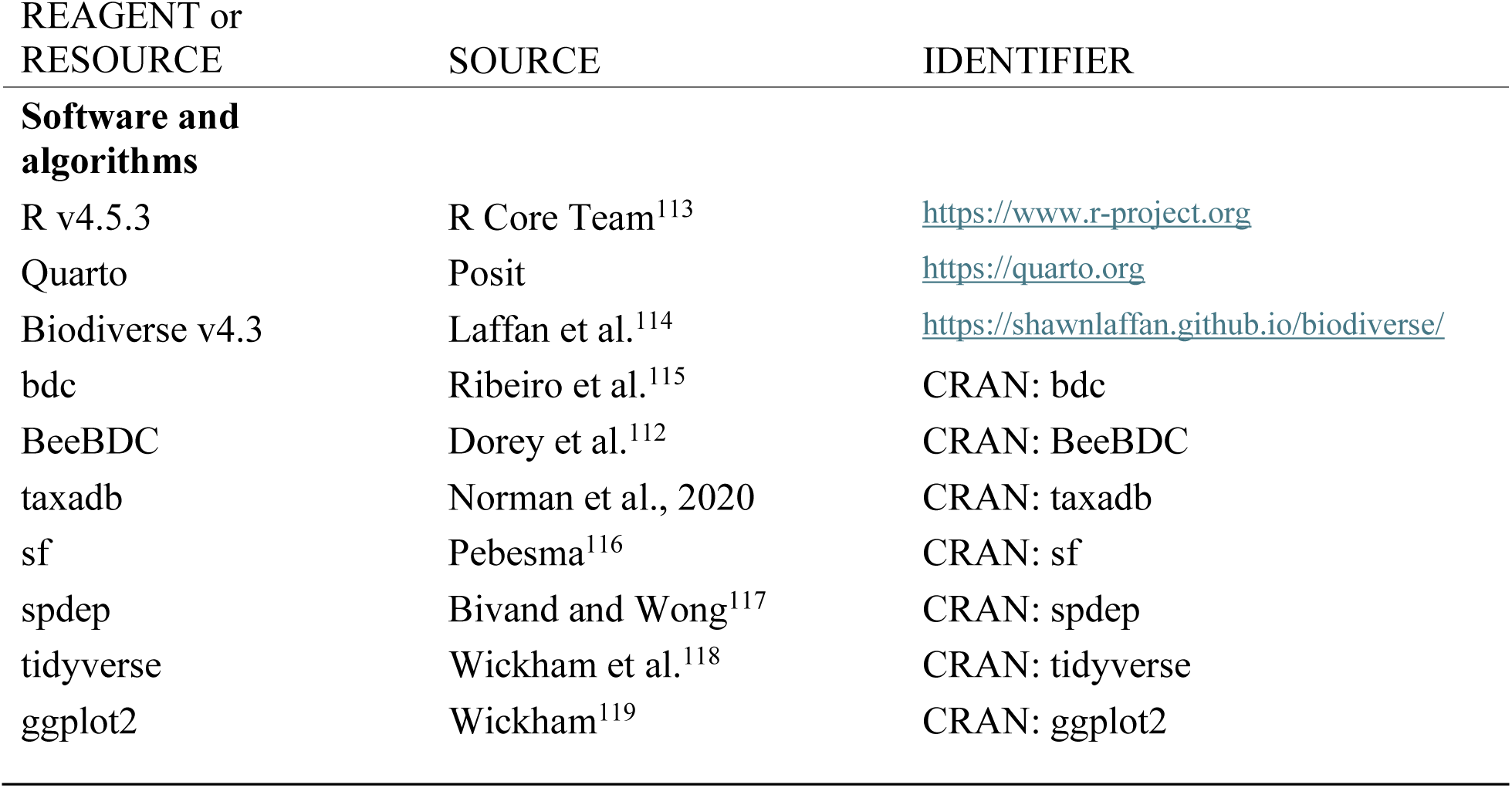

### METHOD DETAILS

#### Bee occurrence data sources

Bee occurrence data for California were assembled from three sources. The first was the Bee Library (library.big-bee.net), an online repository of digitized bee specimen occurrence records compiled by the Big Bee project, a collaboration among 13 universities, research stations, natural history collections, and agencies in the United States^111^, supplied as two occurrence files that were combined before filtering. The second was the United States subset of a publicly available cleaned and standardized global bee occurrence dataset drawing on the Global Biodiversity Information Facility (GBIF), the Symbiota Collections of Arthropods Network (SCAN), Integrated Digitized Biocollections (iDigBio), the United States Geological Survey (USGS), and the Atlas of Living Australia (ALA), together with private collections and records published in the literature^112,120^. The third was the USDA-ARS Pollinating Insects Research Unit, which maintains a database of its own collection together with bees collected in projects for which the unit provided identifications. The three source files contained 2,990,681 raw records in total: 120,516 from USDA-ARS/BBSL, 510,033 from the Bee Library, and 2,360,132 from the Dorey et al. United States subset.

The retained analytical dataset spans specimens collected between 1700 and 2024, with only two records predating 1800 (Figure S2). Records derive from 162 distinct institutional collections.

Institutional codes could be parsed for 513,547 of the 559,021 retained records (91.9%). The largest contributors are the Bee Biology and Systematics Laboratory (BBSL, including specimens from Yosemite National Park, Pinnacles National Park, and other BBSL-identified material), 187,179 records (36.4%); the Essig Museum of Entomology (EMEC), 115,239 (22.4%); the UC Riverside Entomological Collection (UCRC), 41,900 (8.2%); the UC San Diego collection, 28,302 (5.5%); and San Francisco State University (SFSU), 16,274 (3.2%). All percentages are shares of the 513,547 records with parseable attribution rather than of the full dataset. Institutions contributing at least 1% of attributed records are listed in Table S4; a further 148 collections contributed the remaining 49,972 attributed records.

#### Plant occurrence data source

A dataset of 1,383,762 occurrences of native vascular plant species in California, with specimens collected between 1800 and 2017 (Figure S2), was obtained from the Dryad repository (https://doi.org/10.6078/D16K5W)^121^. These data were originally assembled, curated, and validated for a previous study of the native California flora^37^. Because bees are most closely associated with angiosperms, only the angiosperm records were used, comprising 1,333,270 occurrences; these are referred to throughout as the angiosperm occurrence data. No additional cleaning was applied to the angiosperm data, which were used as curated by Baldwin et al.^37^.

#### Data harmonization

All processing was carried out in R v4.5.3^113^. Field names differ among the three source databases, so equivalent columns were mapped to a common set of standardized fields before merging: scientific name, decimal latitude, decimal longitude, catalog number, and a source label, together with the institution, collection, identifier, collector, specimen-number, date, and locality fields used for provenance reporting. All columns were read as text, headers were standardized to lower-case snake_case, internal whitespace was collapsed, and empty strings were converted to missing values. Every record retained a source label identifying the contributing database; this label is carried through each subsequent step and into the final analytical dataset. Raw input files were never modified. Catalog numbers are unique within a collection and, combined with taxonomic and coordinate information, allow specimens duplicated across the three databases to be identified. Catalog numbers were normalized for comparison by squishing whitespace and converting to upper case, so that the same specimen catalogued in two databases with differing spacing or case would match.

The cleaning and validation steps below are implemented in two Quarto documents: 02_create_specimen_datasets.qmd, which performs the initial specimen and name filtering, and 03_data_prep_bees.qmd, which performs all subsequent cleaning. Both write an audit file at every deletion step. Record counts for the full pipeline are given in Table S1.

#### Observation and species-level filtering

Analyses were restricted to specimen-based occurrences rather than opportunistic observations, to reduce the sampling biases documented for community-science platforms^25,122,123^ and to remain consistent with the methodology of Baldwin et al.^37^. Each retained record had to satisfy two conditions.

*Preserved specimen.* The basis-of-record value was normalized by lower-casing and removing spaces and underscores, then compared with “preserved specimen.” Any other value, including a missing value, was treated as not a confirmed specimen and the record was removed. All iNaturalist occurrences and other human-observation records were removed by this rule. The check was not applied to the USDA-ARS/BBSL source, which consists entirely of preserved specimens. To avoid discarding valid specimens carrying a different or missing basis-of-record code, the filter summary records for each source whether the check was applied.

*Species-level name.* A scientific name containing no internal whitespace was treated as a genus-only identification and the record was removed. This check was applied to all three sources, including USDA-ARS/BBSL. Missing names were not caught by this test and were removed at the completeness filtering step below. The rule does not exclude names of the form *Genus* sp. or *Genus* cf. *species*, which contain whitespace; excluding those would require name-level taxonomic evaluation beyond a token count.

Together these rules removed 967,330 of the 2,990,681 raw records (32.3%). The three filtered source files were then merged, giving 2,023,351 standardized records entering the cleaning workflow. Removed records were written to per-source audit files with a plain-language reason for each deletion, and to a single combined audit file.

#### Completeness filtering

Records missing any field required by subsequent cleaning or spatial steps (scientific name, latitude, longitude, or catalog number) were removed before the geographic overlay. Requiring a catalog number means that specimens lacking a catalogued identifier were excluded regardless of the quality of their locality or taxonomic data. This step removed 131,560 records (6.5%), leaving 1,891,791.

#### Geographic validation

Records were restricted to California by a point-in-polygon overlay performed in R with the sf package^116^. The California state boundary shapefile was downloaded from the California Open Data Portal in July 2026^124^, read with sf::st_read(), repaired with sf::st_make_valid(), and transformed to WGS84 (EPSG:4326) to match the coordinate reference system of the incoming records. Occurrence records were converted to point features with sf::st_as_sf() using decimal longitude and latitude in EPSG:4326. A record was retained only if its point intersected the state boundary polygon, evaluated with sf::st_intersects(). Original coordinate columns were preserved alongside the geometry, so no coordinate values were altered. Records falling immediately offshore, or immediately across a state or international border, were removed rather than snapped or buffered inward. Coastal and border records are therefore treated conservatively, and cells along the coastline and the state margin may lose records that fall within the sampled area but outside the boundary polygon.

The overlay retained 723,192 of 1,891,791 records (38.2%) and removed 1,168,599 (61.8%); its effect is summarized in Table S1. The large proportion removed reflects the geographic scope of the source databases rather than data quality, since the Dorey et al. subset is continental in extent and most of its records fall outside California by design.

#### Taxonomic standardization

The bdc package^115^ was used to flag occurrences lacking scientific names or coordinates and coordinates falling outside possible latitude and longitude ranges; flagged records were removed with BeeBDC::summaryFun() and bdc::bdc_filter_out_flags(). No records were removed at this stage, because the completeness filter had already excluded records lacking names or coordinates.

Names were then standardized against two taxonomic authorities in sequence. Verbatim scientific names were first parsed with bdc::bdc_clean_names() and queried against the GBIF taxonomic backbone using bdc::bdc_query_names_taxadb(), with synonyms replaced by accepted names and approximate name suggestions accepted at a string-similarity threshold of 0.85. Parsed and queried names were then harmonized with BeeBDC::harmoniseR() against a global bee taxonomy generated from BeeBDC::beesTaxonomy() and archived as a dated reference file (built 2026-05-07) so that the taxonomic treatment applied here is fixed and reproducible. Because synonym replacement and approximate matching can reassign names, the species total reported below reflects this harmonized treatment rather than the verbatim determinations on specimen labels.

Valid names were matched for 692,883 of the 723,192 California records, leaving 30,309 unmatched. A summary filter applied after harmonization removed 30,491 records in total, retaining 692,701.

#### Duplicate detection and removal

Duplicate records were identified with BeeBDC::dupeSummary()^112^ in two passes. The first compared catalog number and scientific name; the second compared latitude, longitude, and scientific name together with catalog number. Catalog-number matching used minimum thresholds of two characters, five digits, and five digits for numeric-only identifiers, so that short or uninformative identifiers could not generate spurious matches.

Matched records were clustered, and within each cluster the retained copy was selected by ordering records first by source priority — USDA-ARS, then Big Bee, then the Dorey et al. dataset — then by completeness of the latitude, longitude, and scientific-name fields, then by the summary flag column. Because the retained copy is chosen by source priority, the proportion of each database’s records surviving de-duplication reflects that priority rule and is not a measure of data quality. Of the 692,701 records entering de-duplication, 456,042 (65.8%) were unique to a single database, 115,466 (16.7%) were retained copies of specimens present in more than one database, and 121,193 (17.5%) were removed as redundant copies, leaving 571,508.

Cross-database representation was measured by two independent methods. BeeBDC clustering placed 236,659 records in a duplicate cluster, while catalog-number matching independently identified 236,493 records — 34.1% of those carrying both a catalog number and a recognized source label — whose catalog number appeared under more than one source label, an agreement of 99.9% between the two methods. That roughly a third of records appear under more than one source label indicates that these specimen data are broadly accessible through multiple aggregators, and that catalog numbers are recorded consistently enough across them to be matched.

Retention differed markedly among sources, and not only because of the priority rule. The lower proportion of Bee Library records surviving de-duplication reflects duplication within that source: the Bee Library was supplied as two overlapping occurrence files, and redundant copies shared between them were removed irrespective of source priority. Retention rates in Table S2 should therefore be read as a joint consequence of within-source duplication, the source-priority rule, and the subsequent removal of non-native species, rather than as an indication of data quality.

Because de-duplication removed only redundant copies of the same specimen, whether those copies arose across or within source databases, each record in the final dataset corresponds to a single specimen. Throughout this paper, “occurrence record” and “specimen record” therefore refer to the same analytical unit.

#### Non-native species removal

Records of bee species not native to California were removed after de-duplication and before coordinate projection, so that the projected analytical dataset represents the native California bee fauna. Removal was by exact match of the harmonized species name against a fixed list of 30 non-native taxa, comprising introduced species such as *Apis mellifera* and *Megachile rotundata* together with species not known from the western United States, northern Mexico, or southwestern Canada. The list was compiled in advance; for taxa in question, species distributions, collectors, and determiners informed inclusion. Matching was applied to the taxonomically harmonized name rather than the verbatim determination, so that synonyms of listed taxa were also removed.

This step removed 12,487 records (2.2% of the de-duplicated dataset), retaining 559,021. *Apis mellifera* accounted for 11,262 of the removed records. Two listed taxa, *Lasioglossum cressoni* and *Xylocopa appendiculata*, matched no records. The full list and per-taxon counts are given in Table S3.

#### Rare species validation

No abundance threshold was applied at any stage, and species represented by few records were retained, because range-restricted species carry much of the signal in the endemism analysis.

Species represented by five or fewer occurrences statewide underwent a subsequent manual validation step, performed outside the scripted workflow: digitized specimen labels were examined where available, locality information was verified against label data, and California occurrence was confirmed by direct plotting of localities. Records confirmed as legitimate California occurrences were retained, and no records were removed from the analytical dataset on the basis of low species-level occurrence counts.

#### Coordinate system conversion

To match the angiosperm dataset, whose coordinates are in meters under an Albers Conic Equal Area projection, bee coordinates were converted to the NAD83 / California Albers projection (EPSG:3310) using the sf package^116^. This step altered no records. In the final analytical dataset the projected coordinate columns retain the names long and lat for compatibility with earlier versions of the workflow; these columns contain California Albers x and y coordinates in meters, not decimal degrees.

The final analytical dataset contains 559,021 occurrence records representing 1,742 species, 18.7% of the raw records read from the three source files^125^. Record attrition at every step is given in Table S1.

#### Spatial grid construction

Occurrences were mapped onto a uniform grid of 15 × 15 km cells covering California. This grain was chosen as a compromise between spatial resolution and data sufficiency per cell: finer grids would have produced many more empty or sparsely sampled cells, reducing the stability of richness estimates, while coarser grids would have aggregated environmentally distinct areas and obscured spatial variation. The grain also matches Baldwin et al.^37^, permitting direct comparison with the angiosperm data. Because analyses of gridded data can be sensitive to the choice of aggregation unit, the intent was not to identify a universally optimal grain for bees but to use a resolution balancing interpretability, comparability, and record density for this dataset.

Grid cells were generated in Biodiverse v4.3^114^. Grid generation and diversity metric calculation were performed interactively rather than by script, and this forms a documented boundary in the automated reproducibility of the workflow. The Biodiverse output shapefiles are archived with the deposited data so that all downstream analyses can be reproduced without re-running Biodiverse.

### QUANTIFICATION AND STATISTICAL ANALYSIS

#### Sampling adequacy and redundancy

To assess whether apparent richness patterns reflected true diversity or uneven collecting effort, a redundancy index was calculated for each grid cell as 1 − (richness / number of occurrences), which increases as sampling becomes more complete^54^. This indicate whether a region is well sampled (high redundancy) or under-sampled (low redundancy).

Redundancy is undefined for cells containing no occurrence records; such cells were retained in the output and assigned a missing value rather than zero. Cells with a redundancy index below 0.54 were classified as low-redundancy for the purpose of interpreting sampling adequacy, following the threshold used by Garcillán et al.^54^.

Because the number of cells with at least one occurrence differed between bees (1695 cells) and angiosperms (1959 cells) due to the broader extend of sampling of plants, the comparison analyses were run using the cell numbers for bees.

#### Richness estimation and Chao1 comparison

Observed species richness per cell was compared with Chao1-estimated richness to evaluate whether an incidence-based estimator would materially change which cells were identified as relatively low- or high-value. Chao1 estimates richness from the number of species recorded in exactly one and exactly two sampling units, and so uses the incidence of species across records within a cell rather than their abundances. For each cell the richness shortfall (Chao1 minus observed richness) and sampling completeness (observed richness divided by Chao1) were calculated. Overlap between the sets of cells ranked most highly under each metric was then compared at successive percentile thresholds, focusing on the top 10%, 20%, and 30% of cells, and summarized both as the proportion of cells shared and as the Jaccard index.

Because observed richness and Chao1 produced broadly similar rankings, observed richness was retained as the principal metric. Very sparse, low-richness cells were interpreted cautiously, because Chao1 can be unstable when both richness and the number of records are very low. Full diagnostics are given in Figure S1 and Table S7.

#### Diversity and endemism metrics

Native species richness, weighted endemism (WE), and corrected weighted endemism (CWE) were calculated across California in Biodiverse^114^. Species richness is the number of unique native species in a grid cell. WE is the sum of the inverse range sizes of the species present in a cell (1 / number of cells occupied by a species, summed across the species in that cell), which increases the influence of range-restricted species without imposing arbitrary thresholds. CWE standardizes WE by cell richness^80^, reducing the dependence of WE on overall diversity and highlighting cells with disproportionately high concentrations of narrow-ranged taxa^126^. Because CWE corrects for this richness dependence, CWE rather than WE is reported.

#### Hotspot analysis

Spatial clustering of species richness and CWE was assessed with the Getis-Ord Gi* statistic^127,128^, implemented in R with the sf and spdep packages^116,117^. The analysis was run separately for bees and for angiosperms, using the same grid and the same procedure for both. Neighbor relationships among grid cells were defined by queen contiguity, treating cells that share either an edge or a corner as neighbors. Cells were classified as hotspots or coldspots at the 90%, 95%, and 99% confidence levels. A positive z-score with a low p-value indicates significant clustering of high richness or endemism; a negative z-score with a low p-value indicates clustering of low values. All analyses were performed in the California Albers projection.

#### Spatial autocorrelation

Global spatial autocorrelation was calculated using Moran’s I to characterize the spatial scale and intensity of bee diversity patterns. This analysis was applied to the bee data only. Spatial weights were defined by a distance band linking all grid-cell centroids within 22 km, chosen to capture both edge-adjacent (15 km) and diagonally adjacent (approximately 21.2 km) cells on the 15 km grid and thereby approximate queen contiguity. Weights were row-standardized.

Significance was assessed both analytically and by Monte Carlo permutation with 9,999 simulations, using the spdep package. This analysis distinguishes broad, landscape-level spatial structure from fine-scale localized clustering.

#### Concordance between bees and angiosperms

The spatial analyses of richness and CWE described above were applied to the angiosperm dataset as well as to the bee dataset; these analyses were not performed by Baldwin et al.^37^. Following established approaches^38,129–131^, linear regression models were fitted in R to test whether angiosperm richness predicts bee richness and whether angiosperm CWE predicts bee CWE at the grid-cell level.

#### Assessment of sampling bias

Occurrence data assembled from museum and survey collections carry collecting biases that are not distributed evenly in space, and three procedures were used to characterize rather than correct them. First, the redundancy index were calculated for every grid cell, so that cells whose apparent richness rests on thin sampling can be identified and interpreted accordingly. Second, all three source databases were retained through harmonization and de-duplication, with the contributing source recorded on every record, allowing cross-database representation to be quantified directly (see duplicate detection and removal). Third, record counts, deletion reasons, and per-source contributions are reported at every step of the pipeline (Tables S1 and S2), and audit files of removed records can be generated with the deposited data and analysis code, so that the composition of the analytical dataset is inspectable rather than inferred. No post hoc correction, weighting, or spatial thinning was applied to compensate for uneven effort.

